# Distinct waves of ovarian follicles contribute to mouse oocyte production

**DOI:** 10.1101/2024.05.30.596686

**Authors:** Qi Yin, Allan C. Spradling

## Abstract

The earliest growing mouse follicles, wave 1, rapidly develop in the ovarian medulla, while the great majority, wave 2, are stored for later use as resting primordial follicles in the cortex. Wave 1 follicles are known to mostly undergo atresia, a fate sometimes associated with the persistence of steroidogenic theca cells, but this connection is poorly understood. We characterized wave 1 follicle biology using tissue clearing, lineage tracing and scRNAseq to clarify their contributions to offspring and to steroidogenesis. Wave 1 follicles lineage marked by E16.5 *FoxL2* expression in granulosa cells, reach preantral stages containing theca cell layers by 2 weeks. Atresia begins about a week later, during which 80-100% of wave 1 follicles degrade their oocytes, turn over most granulosa cells, but retain theca cells which expand in number together with interstitial gland cells in the medulla. During puberty (5 weeks), these cells ultrastructurally resemble steroidogenic cells and highly express androgen biosynthetic genes. Unexpectedly, the *FoxL2* lineage tag also marked about 400 primordial follicles, located near the medullar-cortical boundary, that become the earliest-activated wave 2 follicles. These “boundary” or “wave 1.5” follicles generate 70-100% of the earliest mature oocytes, while fewer than 26 wave 1 follicles with oocytes survive. Consistent with their largely distinct fates in steroid or oocyte production, granulosa cells of antral wave 1 and wave 2 follicles differentially express multiple genes, including *Wnt4* and *Igfbp5*.

**SIGNIFICANCE STATEMENT:** We studied the ovaries of juvenile mice 2-6 weeks (wk) of age using transcriptomics and cell lineage to characterize the earliest developing groups of follicles. Wave 1 follicles begin growing at birth in the ovarian medulla, but few produce mature oocytes. Instead, most undergo partial atresia and expand steroidogenic cells expressing androgenic genes at puberty. A newly identified primordial follicle subset located at the medulla-cortex boundary are the earliest to activate and grow. Their oocytes give rise to most early offspring, while no more than a few derive from surviving wave 1 follicles. Our results highlight the importance of spatially localized follicle subgroups and show that follicular waves aid reproduction in distinct ways, some of which involve substantial atresia and cellular remodeling.

## INTRODUCTION

Gametes often begin development in subgroups known as “waves” that arise at specific times and gonadal locations (Hirshfield, 1992; Hirshfield and DeSanti, 1995; Gougeon, 1996; Jiménez, 2009; DeFalco and Capel, 2009; Zheng et al., 2014a, b). In mice, the earliest developing follicles, a small group known as “wave 1”, become growing primary follicles in the ovarian medulla by the time of birth without entering dormancy as primordial follicles. The remaining ∼90%, “wave 2”, enter quiescence before postnatal day 5 (P5) by forming primordial follicles within the ovarian cortex to support lifelong fertility (reviews: Edson et al., 2009; McKey et al., 2022) (Figure 1A). In the medulla, wave 1 follicles retain early arising bipotential granulosa cells (BPGs), which express *Foxl2* and *Nr5a2* beginning during early fetal development. In contrast, wave 2 follicles in the cortex acquire epithelial granulosa cells (EPGs), express *Foxl2* only after birth, and most express little *Nr5a2* as primordial follicles (Mork et al., 2012; Zheng et al., 2014a; Niu and Spradling, 2020; Meinsohn et al., 2021a). However, a small subset of non-growing primordial follicles was identified that did express *Nr5a2* and were proposed to be primed for activation (Meinsohn et al., 2021b).

**Figure 1.**
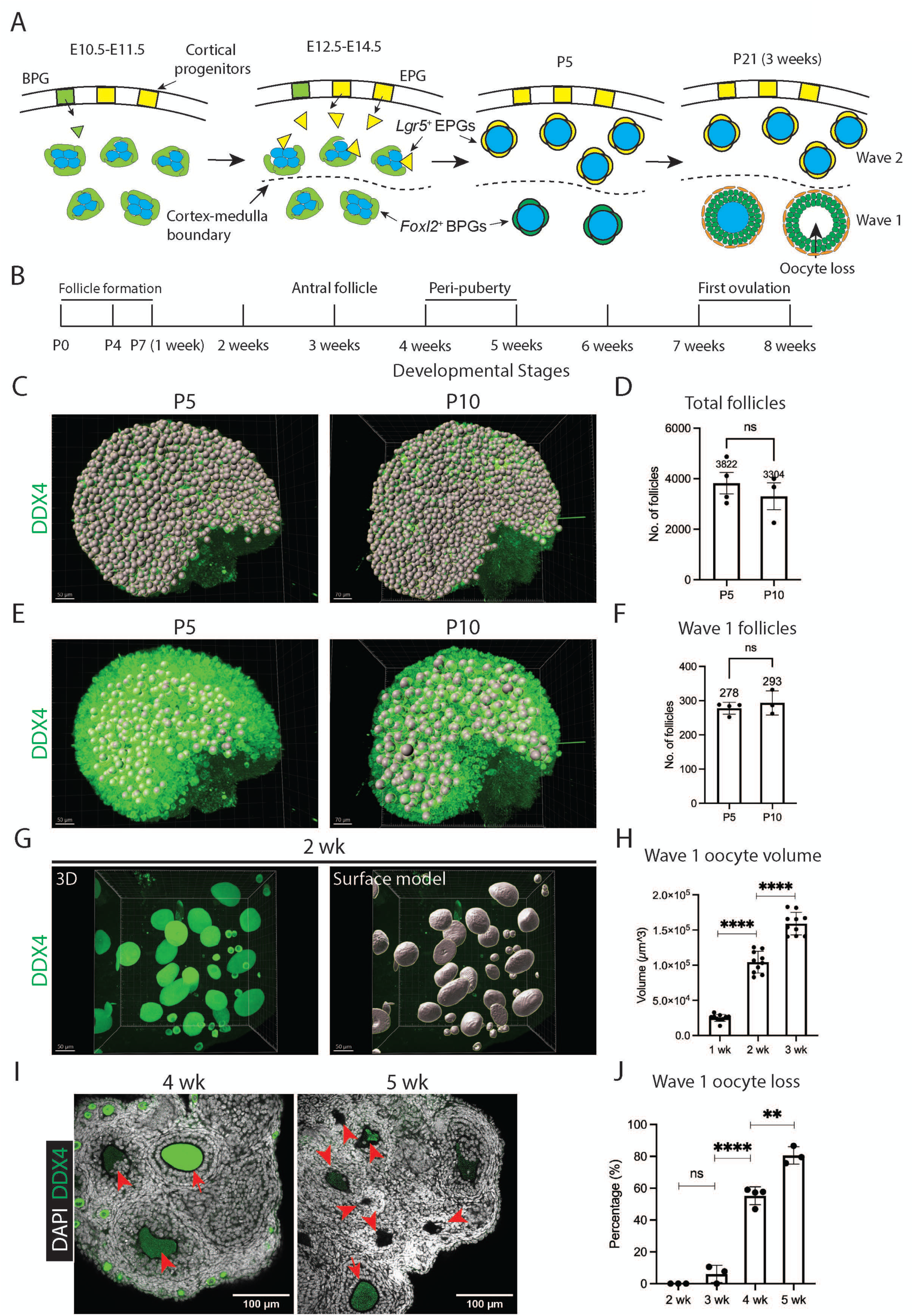
Wave 1 follicles are remodeled prior to peri-puberty. (**A**) Schematic diagrams of fetal and postnatal ovaries at the indicated stages (E = embryonic day; P = postnatal day). Wave 1 follicles, located in the ovarian medulla (below the dashed line), retain bipotential pre-granulosa cells (BPGs, green) and develop without delay. In contrast, cortical follicles, which become quiescent primordial follicles (Wave 2), undergo replacement of BPGs by epithelial pre-granulosa cells (EPGs, yellow). This replacement is completed by P5. By P21 (3 wk), wave 1 follicles undergo atresia, characterized by oocyte loss. Germ cells or oocytes are shown in blue. Developmental stages are indicated in the diagram. Dashed line: Medulla-cortex boundary. (**B**) Postnatal follicular developmental timeline: Wave 1 antral follicles emerge by 3 weeks (Figure S1A-C); Juvenile sexual development (peri-puberty) occurs at 4-5 weeks; Puberty and ovulation take place around 7-8 weeks. (**C**) Annotation of total follicles in P5 and P10 ovaries using Imaris software (gray balls). Each follicles with DDX4-postive oocyte was manually counted and labeled using the Imaris spot model. Scale bars: left, 50 µm; right, 70 µm. (**D**) Total follicle counts in P5 and P10 ovaries (mean ± SD; N = 3-4). ns, not significant. (**E**) Annotation of wave 1 follicles in P5 and P10 ovaries using Imaris software (gray balls). Follicles with oocytes larger than 20 µm in diameter, primarily located in the medulla, are classified as wave 1 follicles at the indicated times. Each gray ball represents one follicle. Scale bars: left, 50 µm; right, 70 µm. (**F**) Wave 1 follicle counts in P5 and P10 ovaries (mean ± SD; N = 3-4). ns, not significant. (**G**) 3D projection of the ovary at P14, reconstructed using the Imaris surface model. Scale bars = 50 µm. (**H**) Oocyte volume increase in wave 1 follicles over time (mean ± SD; N = 10). (Note: Nuclear volume is excluded, as DDX4 is a cytoplasmic protein.). \*\*\*\**p* < 0.0001. (**I**) Representative images of ovary at 4 wk and 5 wk. Left (4 wk): arrowheads indicate an atretic follicle with a distorted oocyte and disorganized granulosa cells; arrow points to a developing healthy secondary follicle. Right (5 wk): arrowheads indicate degrading follicles at various stages; arrow points to a developing healthy secondary follicle. Scale bars = 100 µm. (**J**) Wave 1 follicle remodeling, indicated by oocyte loss over time (mean ± SD; N = 3-4). Oocyte loss begins around 3 wk and continues through 5 wk, accompanied by granulosa cell disorganization and cell death. ns, not significant; ** *p* < 0.01; \*\*\*\**p* < 0.0001. **Figure supplement 1**. Wave 1 follicle development in the early juvenile ovary. **Figure supplement 2**. Wave 1 follicles begin to remodel by 3 weeks.

Once primordial follicles are signaled to grow, their granulosa cells (GCs) start to proliferate and soon form multiple GC layers surrounding the oocyte (Habara et al., 2021). Granulosa cells in secondary follicles send signals encoded by Desert hedgehog (*Dhh*) and Indian hedgehog (*Ihh*) that attract ovarian mesodermal precursors to form outer theca cell layers on follicles and to synthesize steroid hormones (Liu et al., 2015). Additionally, ovarian medullary precursors generate thecal fibroblastic cells and perivascular smooth muscle cells that vascularize the thecal region and enable such follicles to collectively serve as a major systemic steroid hormone source (Guzmán et al., 2023). An ovarian cell type closely related to theca cells, interstitial gland cells, also coordinately form in mice and many other mammals (Jiménez, 2009; Miyabayashi et al., 2015). *Nr5a2* promotes theca cell vascularization that is essential for ovulation (Duggavathi et al., 2008; Bianco et al., 2019; Guzmán et al., 2022). Two classes of steroid hormone-producing Leydig cells, fetal and adult, also arise during testis development (Miyabayashi et al., 2015; Liu et al., 2016).

Follicular cells synthesize specific steroid hormones during their development (Edson et al., 2009; Richards et al., 2017; Niu and Spradling, 2020; Meinsohn et al., 2021a). Thecal steroidogenic cells of late secondary and preantral follicles convert cholesterol to androgens through a series of enzymatic reactions involving steroidogenic acute regulatory protein (StAR), cytochrome P450 family 11 subfamily A member 1 (CYP11A1), hydroxy-delta-5-steroid dehydrogenase, 3 beta- and steroid delta-isomerase 1 (HSD3B1), cytochrome P450 family 17 subfamily A member 1 (CYP17A1), and steroid 5 alpha-reductase 1 (SRD5A1). *Cyp17a1-iCre* (*Cyp17a1* promoter driving codon-improved *Cre*) has been developed and used to label theca cells, the interstitial gland, and some granulosa cells (Bridges et al., 2008). Ovarian androgen production by theca cells during juvenile development is essential for normal maturation and puberty onset (Galas et al., 2012; Walters et al., 2019; Kelava et al., 2022). In addition, thecal androgens diffuse into neighboring GCs, where they are converted into estrogen by the action of hydroxysteroid 17-beta dehydrogenase 1 (HSD17B1) and cytochrome P450 family 19 subfamily A member 1 (CYP19A1). Estrogen produced in this manner signals the hypothalamus to initiate gonadotropin production and is essential for subsequent follicular maturation. After ovulation, the oocyte-free follicle is further modified into a corpus luteum, and its remaining granulosa cells and theca cells undergo luteinization to generate progesterone.

The development and function of wave 1 follicles has remained controversial. Most wave 1 follicles undergo atresia after reaching early antral stages (Byskov et al., 1974; Erickson et al., 1985) during juvenile development (Figure 1B). This suggested they do not primarily serve a direct reproductive function (Hirshfield, 1992; Hsueh et al., 1994; Eppig and Handel, 2012), although these follicles are capable of developing into functional oocytes following superovulation (Lamas et al., 2021) and could support fertility in an irradiated rat model (Guigon et al., 2003). Theca cells of atretic follicles often survive and persist in the ovary (Erickson et al., 1985; Magoffin, 2005), suggesting a function in steroid production. However, when *Foxl2-CreERT2* was used to lineage label the granulosa cells of wave 1 follicles, marked follicles substantially contributed to offspring during the initial months of fertility (Zheng et al., 2014b).

Here we studied wave 1 follicle development and gene expression in detail to gain deeper insights into follicular waves. We found that lineage labeling using *Foxl2-CreERT2* activated in early fetal stages is not specific for wave 1 follicles. Granulosa cells in about 400 primordial follicles are also marked near the medulla-cortex boundary in 2week old mice, hence we termed them “boundary” or “wave 1.5” follicles. Marked boundary follicles produce more than 70% of early fertile oocytes, while no more than 26 intact wave 1 follicles survive to puberty. Boundary follicles probably correspond to the resting primordial follicle population expressing *Nr5a2* described by Meinsohn et al. (2021a, b). Thus, most of the *FoxL2-CreERT2* labeled follicles contributing to offspring observed by Zheng et al. 2014a likely derived from boundary follicles rather than wave 1 follicles.

Ovarian developmental and gene expression studies during 2-6 wks provided molecular insight into gamete waves and follicular atresia. Wave 1 follicular oocytes and granulosa cells mostly turn over beginning at about 3 wk. However, their theca cells increase in number and join interstitial gland cells to form by puberty a mass of active steroidogenic medullar cells that retain morphological evidence of their origin from atretic follicles. These steroidogenic cells highly express androgenic genes when granulosa cells are still reduced in number by atresia. The granulosa cell gene expression in developing wave 2 follicles also shows differences from that of wave 1 follicles, indicating that the distinct developmental fates of wave 1 and wave 2 follicles may depend on intrinsic gene expression differences in addition to differing levels of gonadotropins.

## RESULTS

### Wave 1 follicle development

Advances that allow tissue clearing (Faire et al., 2015; Feng et al., 2017; Fiorentino et al., 2020; McKey et al., 2022), now make it possible to image an entire mouse ovary without sectioning. To better understand follicle waves, we employed whole mount staining combined with clearing-enhanced 3D (Ce3D) method to analyze wild type C57BL/6J ovaries at the cellular level (Li et al., 2017). Multiple ovaries were reconstructed during weeks (wk) 1-5 of development (Table S1). In C57BL/6J mice, puberty typically occurs between 4 and 5 wk of age, but the first ovulation, which signals sexual maturity, doesn’t take place until 7 wk (Figure 1B) (Hogan et al., 1994; Nelson et al., 1990; Zhou et al., 2007).

These studies allowed the number of wave 1 and wave 2 follicles to be accurately determined. Wave 1 follicles begin to grow into primary follicles around the time of birth (Figure S1A, B), as confirmed by gene expression analysis (Niu and Spradling, 2020). Following whole-ovary imaging using a confocal microscope, the samples were analyzed using Imaris software. Since nearly all oocytes are enclosed by granulosa cells by P5, it becomes feasible to accurately quantify the number of follicles. At P5 and P10, we counted approximately 3,822 and 3,304 total follicles, respectively (Figure 1C, D), and 278 and 293 wave 1 follicles (Figure 1E, F). Despite variations in total follicle numbers among samples of the same age, the number of wave 1 follicles remained relatively consistent.

By 2 wk, wave 1 follicles in the medulla had developed into secondary follicles containing 2-4 layers of granulosa cells and 1–2 layers of theca cells (Figure S1C). At 3 wk, some follicles further developed into antral follicles with multiple granulosa cell layers, within which granulosa cell apoptosis was observed (Figure S1D). Using the Imaris software surface modeling, we quantified oocyte volume from 1 wk to 3 wk and observed a significant increase during this period (Figure 1G, H).

### Wave 1 follicles undergo partial atresia with theca cell survival

Starting around 3 wk, rather than continuing to develop, wave 1 follicles begin to undergo atresia (Table S1; Supplemental Movies). We investigated the kinetics, cell specificity and completeness of cellular turnover from 3 wk to 5 wk. Atretic follicles initially showed oocyte distortion followed by loss, initially leaving behind disorganized granulosa within the basement membrane but without affecting the theca cell layers (Figure 1I). Atretic follicles progressively decreased in size and developed a cavity beneath the granulosa layer where the oocyte had previously resided. Oocyte loss from wave 1 follicles increased significantly between 3 and 4 wk, exceeding 80% by 5 wk. (Figure 1J). Electron microscopy revealed that during atresia, as oocytes degenerate, microvilli from both oocytes and granulosa cells retract from the zona pellucida (ZP). The shrinking of the cavity is marked by folding of the ZP, and in atretic follicles, remnants of the ZP appear to prevent further collapse of the cavity structure (Figure S2A). ZP remnants in the follicular cavity have previously been used as a morphological indicator of atretic follicles (Myers et al., 2004). We confirmed that these events also occur in the DBA/2J strain, where atretic follicles are also first observed around 3 wk. By 7 wk, corpora lutea from ovulated oocytes and cavities left behind by wave 1 oocyte loss were evident (Figure S2B). Thus, by combining whole mount staining with Ce3D clearing method, we found that the great majority of wave 1 follicles lose their oocytes prior to puberty and do not contribute to early ovulation.

The loss of oocytes from most wave 1 follicles is not simply due to atresia of follicles whose oocytes are defective, as superovulation at these stages can rescue many of them (Lamas et al., 2021). Etymologically, atresia originates from the Greek “a-” (meaning “no”) and “tresis” (meaning “perforation”), referring to the absence or closure of an opening, or cavity. Atresia is sometimes used simply to describe the complete turnover of follicles, which was not observed for wave 1 follicles. We prefer the terms ‘partial atresia’ or ‘remodeled follicles’ to emphasize the potential function of those structures that remain (Hsueh et al., 1994).

To analyze the cell specificity of wave 1 partial atresia, we exploited *Foxl2-CreERT2* mouse line and *R26R-LSL-EYFP* reporter line to selectively lineage label GCs associated with wave 1 follicles (Figure 2A). Similar genetic models have previously been used to study wave 1 follicle development and the duration of their fertility contribution based on tissue sections (Mork et al., 2012; Zheng et al., 2014a). Following brief administration of Tamoxifen (TAM) at embryonic day 16.5 (E16.5), we observed a distinct labeling pattern along the medulla–cortex boundary. Most wave 1 follicles in the medulla showed mosaic EYFP-labeling of GCs at 2 wk (due to the incomplete efficiency of cre-lox recombination) while GCs in the largely cortical wave two follicles remained unlabeled (Figure 2B, C). On average, 201 preantral wave 1 derivatives—representing approximately 90% of wave 1 follicles—exhibited mosaic labeling in granulosa cells (Figure 2E, F). By 5 wk, after extensive wave 1 atresia and remodeling, an average of only 26 antral follicles containing healthy oocytes and EYFP-mosaic GC layers were observed, approximately 70% of all antral follicles (Figure 2D-F).

**Figure 2.**
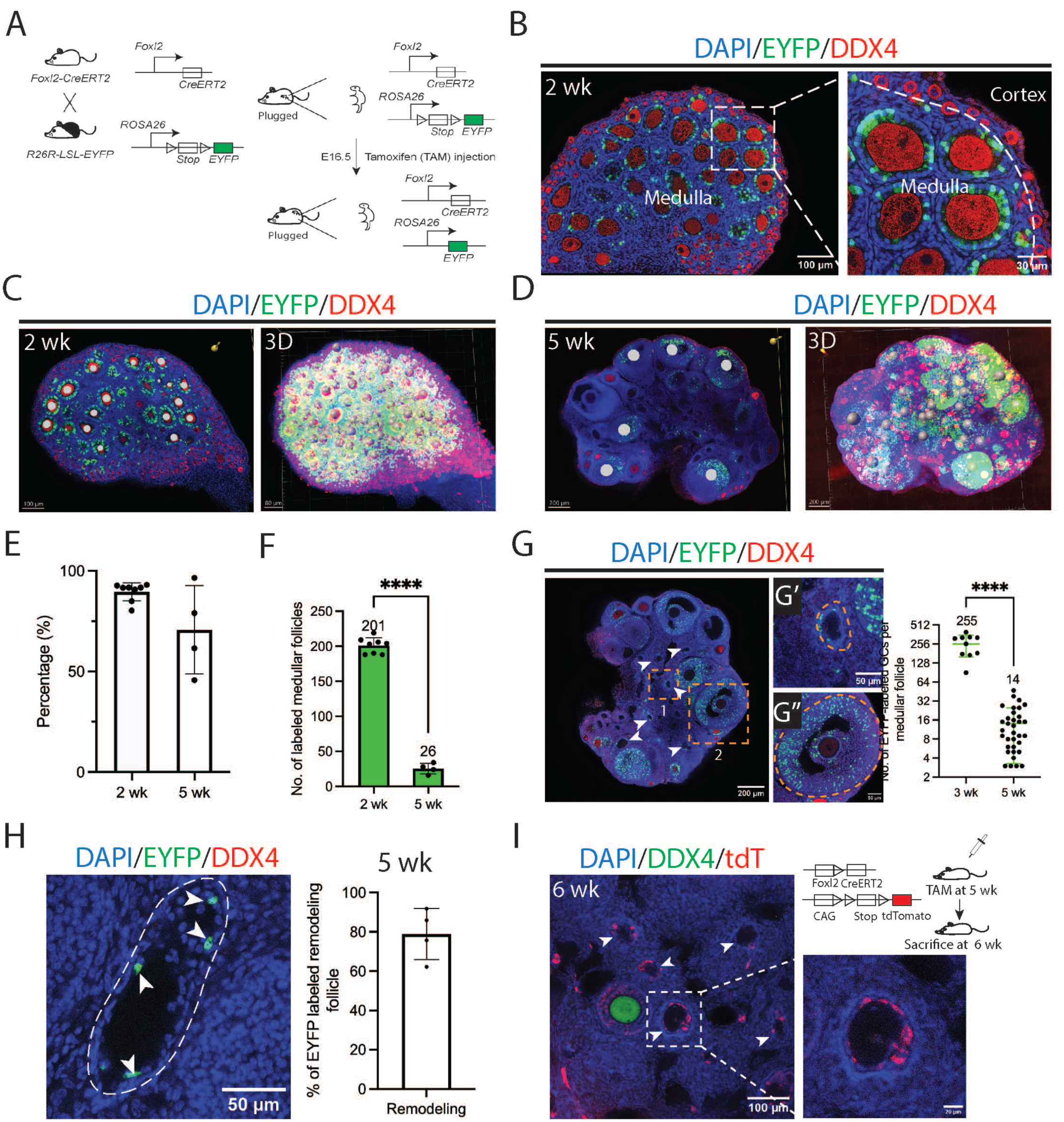
Wave 1 follicles lose granulosa cells during remodeling. (**A**) Schematic diagram illustrating the activation of *Foxl2*-driven EYFP expression following TAM injection at E16.5. TAM administration at E16.5 induces the excision of a stop cassette, leading to EYFP expression in granulosa cells of follicles (see Methods). (**B**) Efficient EYFP labeling of granulosa cells in wave 1 medullary follicles of a 2-week ovary following TAM injection at E16.5, as shown in (A), magnified in the right. Scale bars: left, 100 µm; right, 30 µm. (**C**) EYFP-labeled follicles that have reached the primary/secondary follicle stage by 2 weeks are quantified using Imaris software, with annotation performed using the spot model. Left: A section showing annotated labeled follicles (white circles). Right: 3D projection using Imaris software, where labeled follicles are represented by gray spheres. Scale bars: left, 100 µm; right, 80 µm. (**D**) By 5 wk, EYFP-labeled follicles that have reached the preantral/antral follicle stage are quantified using Imaris software and annotated with the spot function. Left: A section showing annotated labeled follicles (white circles). Right: 3D projection using Imaris software, with labeled follicles represented by gray spheres. Scale bars = 200 µm. (**E**) Over 90% of wave 1 follicles contained EYFP-positive granulosa cells at 2 wk. At 5 wk, approximately 70% of normal antral follicles contained EYFP-positive granulosa cells. (**F**) Quantification of wave 1 follicle numbers (representative images shown in C-D) shows a decline from 2 to 5 weeks. \*\*\*\**p* < 0.0001. Each dot represents one sample. (**G**) Representative section from a 5-week ovary reveals ongoing remodeling of wave 1 follicles. Follicles near the medullary core appear small and without oocytes (arrowheads), with few remaining EYFP-labeled granulosa cells, as seen in the follicle boxed in “1” and enlarged in G’. In contrast, a normal antral follicle in rectangle “2” enlarged in G’’, contains numerous EYFP-labeled granulosa cells. Right: The average number of EYFP-labeled granulosa cells in medullary follicles before remodeling (3 weeks) is compared to remodeled medullary follicles at 5 weeks. Each point represents one analyzed follicle. \*\*\*\**p* < 0.0001. Scale bars: left, 200 µm; right, 50 µm. (**H**) About 80% of remodeled follicles still retain some EYFP-labeled granulosa cells at 5 wk, consistent with their wave 1 origin. Dashed line: follicle boundary; arrowheads: EYFP-positive granulosa cells. Each point represents one analyzed follicle. (**I**) Remnant granulosa cells of remodeled wave 1 follicles continue to express Foxl2 at 5 weeks. TAM was injected into Foxl2-CreERT2; R26R-LSL-tdTomato mice at 5 weeks, and ovaries were collected one week later. Arrowheads indicate remodeled follicles, with one remodeled follicle (outlined in the rectangle) enlarged on the right. Scale bars: left, 100 µm; right, 20 µm. **Figure supplement 3**. Lineage labeling using Foxl2-CreERT2 and R26R-LSL-tdTomato

By comparing the number of labeled GCs in healthy follicles at 3 wk with those in remodeled follicles at 5 wk, we found that remodeled follicles lost not only oocytes but also 94% of their GCs (Figure 2G). The remaining GCs were located adjacent to the cavity left by the oocyte and approximately 80% of remodeled follicles stilled contained labeled GCs (Figure 2H). To ask whether these remnant GCs remained active, lineage labeling was carried out at 5 wk and analyzed one week later. The results showed that many GCs in remodeled follicles still remain intact adjacent to holes left by oocytes and express *Foxl2* (Figure 2I). The turnover of wave 1 follicles was also observed using a different reporter mouse line *R26R-LSL-tdTomato* (Figure S3A-C).

### Identification of an early activating subpopulation of primordial follicles

Because only about 26 normal-appearing wave 1 follicles with oocytes survived even to 5 wk, which seemed too few to support the months of early fertility ascribed to wave 1 by previous reports (Zheng et al., 2014a), we looked for follicles other than wave 1 that might be also labeled by *Foxl2* expression. We observed that *Foxl2-CreERT2* activated by TAM injection at E16.5 in 2 wk juvenile ovaries also labels a subset of primordial follicles located near the medulla-cortex boundary (Figures 3A). A mean of 421 labeled “boundary” follicles mosaic for 1-4 EYFP-positive GCs were identified among 5,843 total follicles (7%) (Figure 3B). Because only 1-4 GCs where labeled from among an average of 8 GCs (Chen et al., 2022), some additional boundary follicles were likely missed for stochastic reasons due to the low efficiency of cre-lox recombination. Applying the Poisson distribution based on the relative proportion of follicles with 1 or 2 labeled GCs, suggested that 90% of boundary follicles were detected.

**Figure 3.**
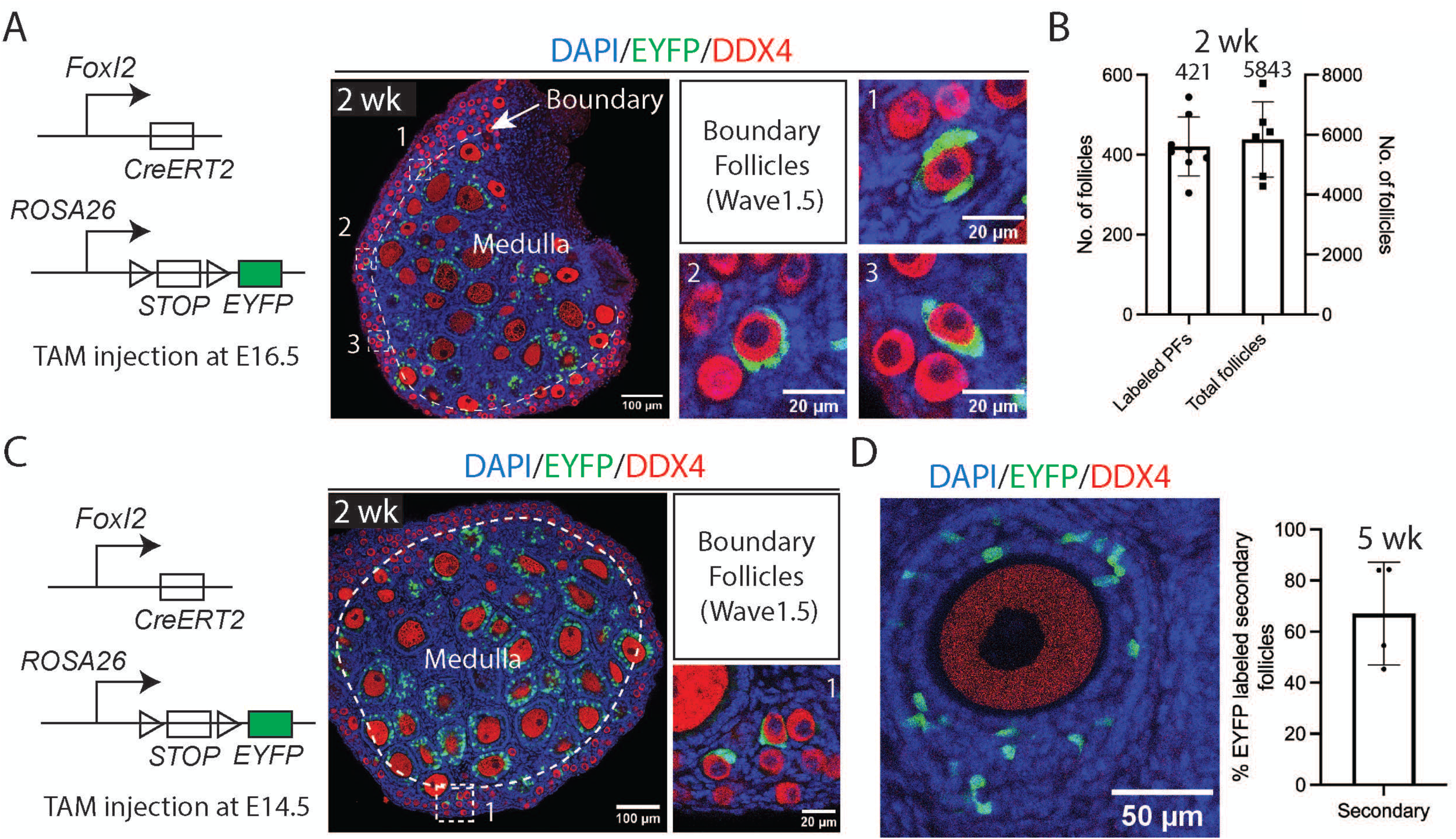
The labeling and quantification of boundary follicles. (**A**) At 2 wk, one or more granulosa cells in some primordial follicles located at the medulla-cortex boundary are labeled with EYFP (enlarged in “1”, “2”, “3”) following TAM injection, as shown in the schematic. These follicles are referred to as boundary follicles or, in accordance with the wave classification, wave 1.5 follicles. Dash line, medulla-cortex boundary. Scale bars: left, 100 µm; right, 20 µm. (**B**) Quantification shows that 7% (421 ± 74) of the total 5,843 ± 1,256 primordial follicles contained EYFP-labeled granulosa cells in the labeled experiment shown in (A). Each point represents one analyzed sample. Labeled PFs: Labeled primordial follicles. (**C**) Representative images of *Foxl2-CreERT2; R26R-LSL-EYFP* mouse ovary injected with TAM at E14.5. Boundary follicles labeled with EYFP are enlarged on the right. Dash line indicates the medulla-cortex boundary. Scale bars: left, 100 µm; right, 20 µm. (**D**) Approximately 70% of secondary follicles in 5 wk ovaries contain EYFP-labeled granulosa cells, suggesting they originate from labeled primordial follicles (wave 1.5) at 2 wk. TAM was injected at E16.5. Each point represents one analyzed sample. Scale bar = 50 µm.

Because of the biological interest of this subgroup of primordial follicles, we repeated activation of *Foxl2-CreERT2* at E16.5 in a separate experiment with a different reporter line, *R26R-LSL-tdTomato.* The results confirmed the existence and properties of boundary follicles (Figure S3A-D). Boundary follicles also identified following TAM injection at E14.5 although at a lower frequency (Figure 3C). Thus, the existence of this subclass was confirmed in multiple experiments.

Primordial follicles begin to activate after 2wk (Zheng et al. 2014a). If boundary follicles behave like other primordial follicles, we would expect subsequently only 7% of recently activated wave 2 growing follicles would show EYFP mosaic GCs at later times. Instead, we found that among secondary follicles at 5 wk, nearly 70% contained EYFP-positive GCs (Figure 3D), indicating that boundary follicles activate at the time of the earliest known primordial follicles, and rise from 7% of quiescent to >70% of early activating follicles imply they comprise most of the earliest activated primordial follicles. Moreover, they correspond in location and properties to the inactive but poised primordial follicles described by Meinsohn et al. (2021a, b).

### Partial atresia of wave 1 follicles does not affect theca cells which expand in number

We then used lineage labeling to investigate the fate of wave 1 theca cells after the onset of atresia. *Cyp17a1* is an essential gene involved in steroidogenesis and is expressed in the steroidogenic theca cells (STCs) and interstitial gland cells (IGCs). We validated that theca cells strongly express the steroid biosynthetic enzyme CYP17A1 from 3 wk to 5 wk using immunofluorescent staining (Figure S4A). To label STCs, we crossed the *Cyp17a1-icre* transgenic mouse line with the *R26R-LSL-tdTomato* reporter line (Figure 4A). In 2 wk ovaries, tdTomato expression was observed in theca cells sheathing large medullar follicles (Figure 4B; arrowhead). Additional cells intertwined with theca cells, which appear to be IGCs (Figure 4B; boxed region), were visualized using Imaris and pseudo-colored using the surface model to highlight their structure (Figure 4C). A relatively small number of GCs also become labeled initially (Figure S4B, arrowheads), possibly because GC precursor cells briefly express *Cyp17a1* during embryonic stages (Mawaribuchi et al., 2014).

**Figure 4.**
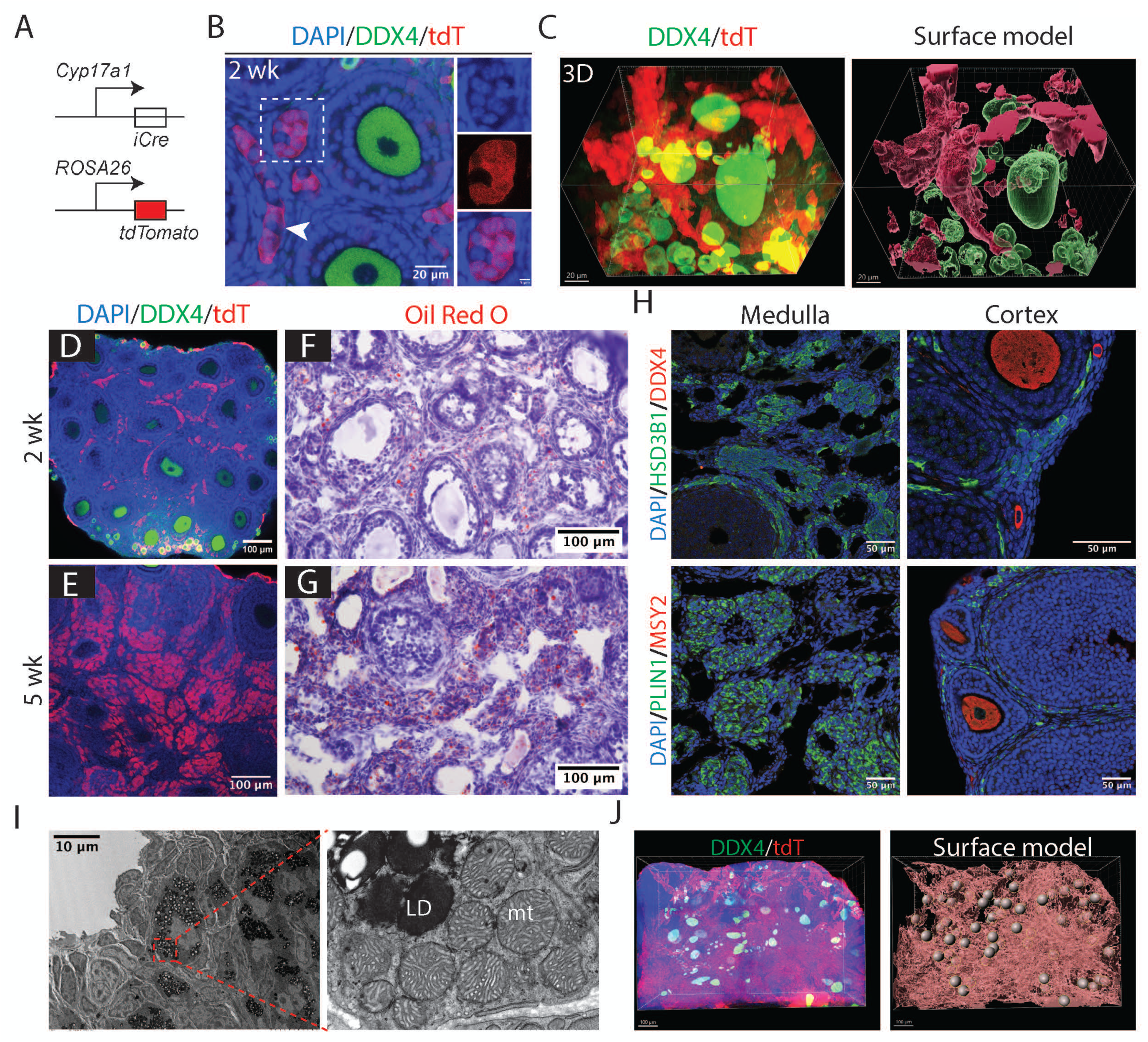
Follicles remodel to form an interconnected stromal network rich in thecal secretory cells. (**A**) Schematic diagram illustrating lineage labeling of *Cyp17a1*-expressing theca cells, interstitial gland cells, and potentially other cell types using *Cyp17a1*-iCre; *R26R-LSL-tdTomato* mouse model. (**B**) 2 wk ovary showing tdTomato-labeled theca cells sheathing follicles (arrowhead) and putative interstitial gland cells between follicles (rectangle). Scale bars: left, 20 µm; right, 5 µm. (**C**) 3D reconstruction of follicles with surrounding labeled theca cells and interconnected interstitial gland cells in a 2 wk ovary. Left: 3D projection using Imaris software. Right: Pseudo-color annotation using the Imaris surface model. Green: Oocytes. Red: *Cyp17a1*-expressing cells, including theca and interstitial gland cells. Scale bars = 20 µm. (**D-E**) Representative images of 2wk (D) and 5 wk (E) ovaries showing a dramatic increase in labeled theca, interstitial, and possibly other Cyp17a1-lineage cells in the ovarian medulla. Scale bars = 100 µm. (**F-G**) Oil Red O staining (red) reveals a significant increase in lipid droplet storage in ovarian medullary cells from 2 wk (F) to 5 wk (G) ovaries. Scale bars = 100 µm. (**H**) Immunofluorescence staining of HSD3B1 and PLIN1 at 5 wk shows that genes required for steroidogenesis and lipid droplet storage are highly expressed in medullary cells. Scale bars = 50 µm. (**I**) Electron microscopy of a remodeled follicle reveals cells with abundant lipid droplets (LD) and characteristic mitochondria (mt) (boxed region magnified in right panel). Scale bar = 10 µm. (**J**) 3D reconstruction of a 5 wk ovary labeled by tdTomato expression activated by *Cyp17a1-iCre*. Left: 3D projection using Imaris software. Right: Pseudo-color annotation using the Imaris surface model. Red: Thecal/interstitial network structure. Balls: Oocyte locations. Scale bars = 100 µm. **Figure supplement 4**. Expansion of theca cells and its related cells - interstitial gland population.

Between 2 wk to 5 wk, the number of tdTomato-labeled cells associated with nuclear cavities dramatically increases as wave 1 follicle atresia and remodeling proceeds to involve more such follicles (Figure 4D, E). Because over 90% of GCs turn over as part of wave 1 remodeling (Figure 2G), the tdTomato-positive cells associated with remodeled wave 1 follicles by 5 wk mostly represent abundant STCs and IGCs. Oil red O staining documented abundant lipid droplet accumulation in these cells from 2 wk to 5 wk (Figure 4F, G). Antibody staining revealed abundant expression of the steroidogenic enzyme HSD3B1 and the lipid storage protein PLIN1 in these cells, which are located in the medulla (Figure 4H). Electron microscopy confirmed that expanded wave 1 follicular cells display distinctive lipid droplets and mitochondria characteristic of STCs and IGCs (Figure 4I, S4C). Using Imaris software surface model, we reconstructed labeled STCs and IGCs throughout the entire ovary. By 5 wk, atretic wave 1 follicles had generated a cellular cluster within the medulla that was dominated by expanded numbers of steroidogenic cells (Figure 4J). This structure persists within the ovary at least until 7 wk (Figure S4D).

### Mapping cell transcription during wave 1 remodeling

To gain further insight into wave 1 follicle development and remodeling, we analyzed gene expression in single cells from ovaries every wk from wk2 to wk6, covering pre- and peri-pubertal stages (Figure 5A; Supplemental Datasets S1-S6). A total of 31 initial clusters (c0–c30) were identified, with nearly all samples contributing to each cluster (Figure 5B). Nine major cell groups were identified (Figure 5C), based on their expression of marker genes (Figure 5D) (Niu and Spradling, 2020; Morris et al., 2022). These comprised 10 granulosa cell clusters, 8 stromal cell clusters, as well as theca cells, hematopoietic cells, endothelial cells, pericytes, smooth muscle cells, and epithelial cells. Four clusters with low UMIs (c6, c11, c15, and c29) were excluded from subsequent analysis (Figure S5A). Additionally, cluster c14 was removed due to potential doublets, as it overlapped with several other clusters; however, we cautiously named it ‘to be determined’ (Figure S5B). Notably, all known ovarian somatic cell types were identified within one or more clusters, but germ cells probably were too rare in these juvenile ovaries to form a cluster, although ovarian germ cells have been extensively studied at earlier times by these methods (Niu and Spradling, 2022) or similar stages using different methods (Morris et al., 2022; Isola et al., 2024).

**Figure 5.**
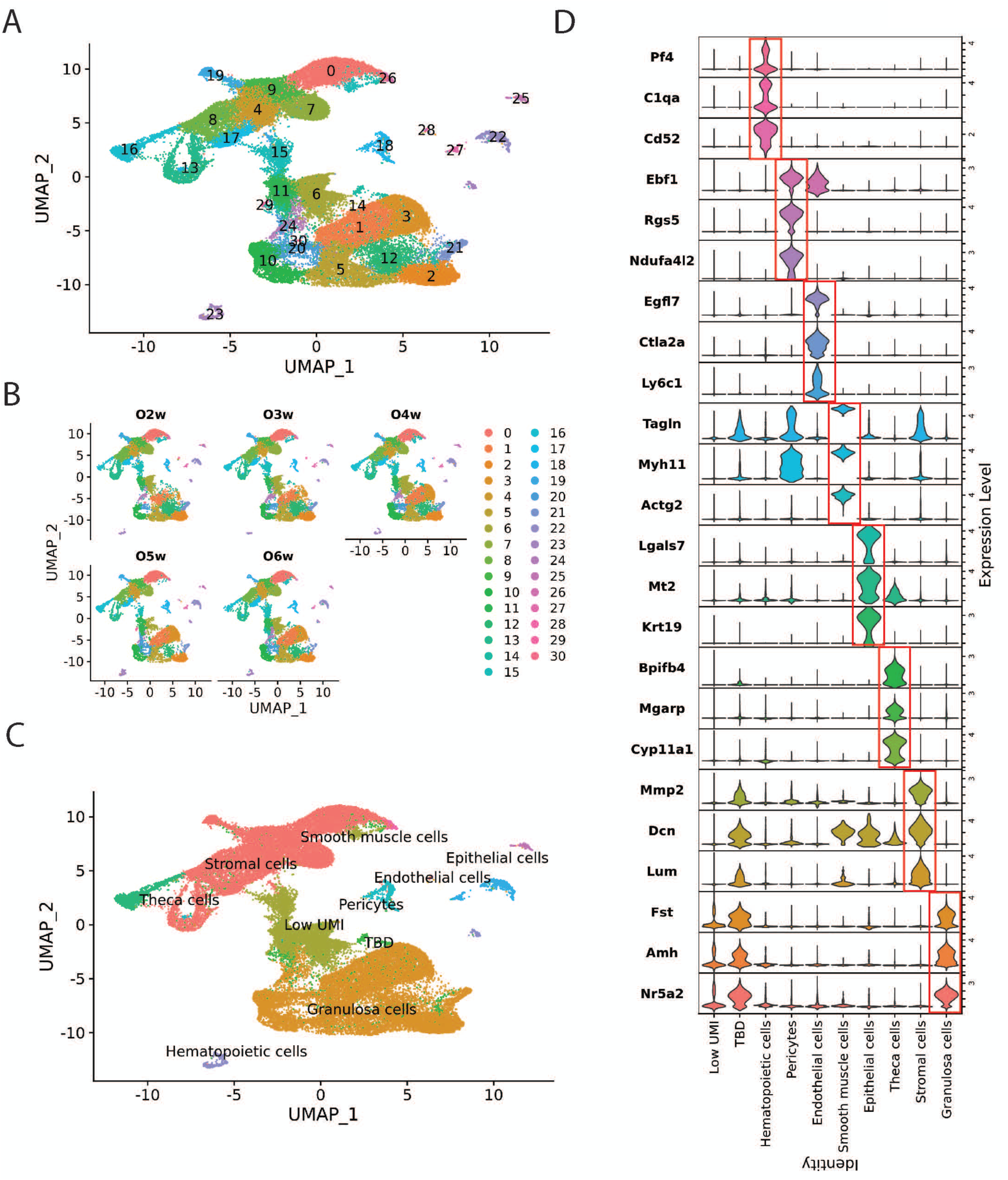
Single-cell RNA sequencing of 2 wk-6 wk ovaries. (**A**) UMAP plot of ovaries from 2 wk to 6 wk, showing 31 initial clusters (c0-c30) represented in different colors. Sequencing data from different ages were merged before summit to Seurat. (detailed information in the methods section). (**B**) The ovarian cells from 2 wk to 6 wk contribute to almost every cluster in the UMAP plot. O2w: scRNA sequencing data from 2 wk ovary. O3w: 3 wk ovary. O4w: 4 wk ovary. O5w: 5 wk ovary. O6w: 6 wk ovary. (**C**) The cell type groups are indicated by colored regions, with their relationship to the numbered initial clusters as shown in A. Granulosa cells: c1, c2, c3, c5, c10, c12, c20, c21, c24 and c30; Stromal cells: c0, c4, c7, c8, c9, 13, c17 and c19; Theca cells: c16; Hematopoietic cells: c23; Endothelial cells: c22 and c27; Pericytes: c18; Smooth muscle cells: c26; Epithelial cells: c25 and c28; Low UMI: c6, c11, c15 and c29; TBD (To be determined): c14. (**D**) Multiviolin plot of selected marker gene expression across different cell types. Y-axis: Gene names with expression levels normalized for display. X-axis: Cell types.

The initial clustering further validated the picture of wave 1 development obtained from lineage studies. Cluster c1 expressed wave 1 BPG granulosa cell markers like *Nr5a2*, *Hsd3b1*, and *Inha*, *Inhba* and increased 4-fold in cell number between wk2-wk4, before falling at wk 5 as wave 1 granulosa cells turn over (Supplemental Dataset S6). Cluster c16 highly expressed theca cell steroidogenic genes such as *Cyp11a1*, *Cy17a1*, and *Plin1*. The number of c16 cells peaked at puberty (wk5). The outer “theca externa” is enriched in angiogenic cells, blood vessels, and fibroblastic cells. Clusters corresponding to endothelial cells expressing *Flt1* (c22) and blood cells expressing *Laptm5* (c23), increased nearly 4-fold between wk3 and wk5, consistent with such vascularization.

### Analyzing GC gene expression

We re-clustered the 10 initial GC clusters in Seurat, yielding 18 new clusters gc0-gc17 (Figure 6A) and classified them into groups corresponding to their likely association with primordial (gc11), primary (gc3, gc8), secondary (gc1, gc4, gc9), antral (gc2, gc7, gc10, gc13), and remodeling (gc0, gc6) follicles, as well as mitotically cycling cells(gc5, gc12, gc14, gc15, gc16, and gc17) (Figure 6B, C). GCs from all follicle stages were identified across all samples (Figure S5C). Single cell pseudotime trajectory analysis using Monocle3 identified distinct developmental paths from primordial follicles to either antral follicles or remodeled follicles (Figure 6D). Once primordial follicles reached the secondary stage, some progressed along path 1 toward antral follicles, while others followed path 2 toward remodeled follicles.

**Figure 6.**
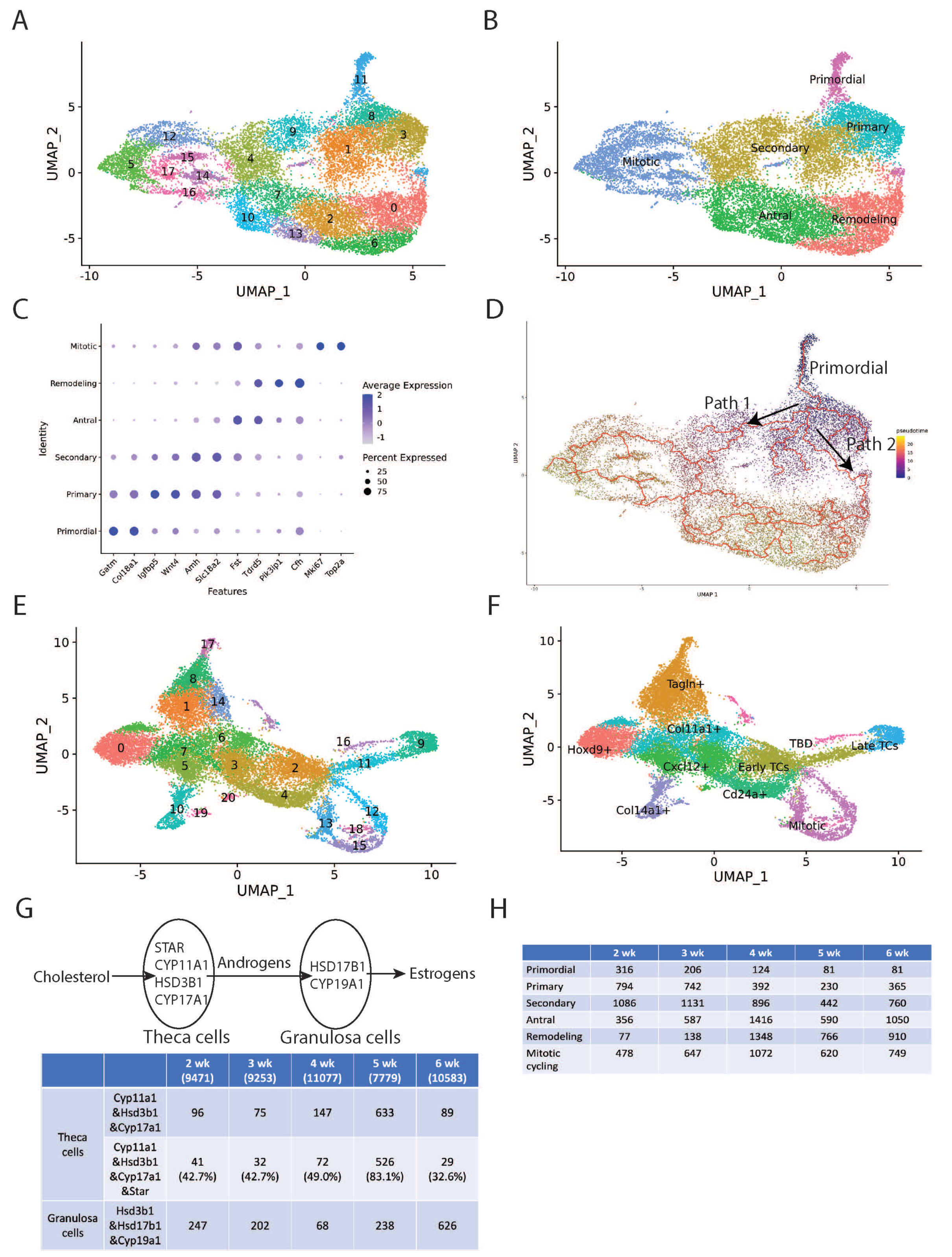
Further re-cluster analysis of sequencing data. (**A**) UMAP plot of re-clustered granulosa cells from Figure 5A, showing 18 clusters (gc0-gc17). (**B**) The deduced ovarian follicle developmental stages are labeled and indicated by colored regions with their relationship to the numbered clusters shown in A. Primordial: gc11; Primary: gc3 and gc8; Secondary: gc1, gc4 and gc9; Antral: gc2, gc7, gc10 and gc13; Remodeling: gc0 and gc6; Mitotic: gc5, gc12, gc14, gc15, gc16 and gc17. (**C**) Dot plot of marker gene expression in granulosa cells across different follicle stages. Y-axis: Cell types. X-axis: Gene names with expression levels normalized for display. (**D**) Pseudotime trajectory analysis using Monocle3 reveals two pathway in secondary follicles: Path 1 leads to further development into antral follicles, while Path 2 results in remodeling follicles. (**E**) UMAP plot of re-clustered theca cell and stromal cells Figure 5A, showing 21 clusters (g0-g20). (**F**) The deduced mesodermal cell subgroups are labeled and indicated by colored regions and their relationship to the numbered clusters shown in panel E. Hoxd9+: g0; Tagln+: g1, g8, g14 and g17; Early TCs: g2 and g11; Late TCs: g9; Cxcl12+: g3, g5 and g20; Cd24a+: g4; Col11a1+: g6 and g7; Col14a1+: g10 and g19; Mitotic: g12, g13, g15 and g18; TBD (To be determined): g16. (G) Diagram of steroidogenic gene expression in theca and granulosa cells and their role in androgen and estrogen synthesis. Top panel: The diagram illustrates the synthesize of androgen in theca cells and estrogen in granulosa cells from cholesterol. Bottom panel: The estimated numbers of steroidogenic theca cells expressing three key steroidogenesis genes in the sequencing data— *Cyp11a1*, *Hsd3b1*, and *Cyp17a1*—are shown, along with those expressing all four essential genes (*Cyp11a1*, *Hsd3b1*, *Cyp17a1*, and *Star*). The total cell number are shown in the first row. From 2 wk to 5 wk, the theca cell population increases, followed by a decline at 6 weeks. Meanwhile, the number and percentage of theca cells expressing all four essential genes progressively increase. In contrast, the estimated numbers of granulosa cells expressing *Hsd3b1*, *Hsd17b1*, and *Cyp19a1* follow a fluctuating pattern, decreasing from 2wk to 4 wk before rising again at 5 wk. Notably, 2 wk corresponds to the time of minipuberty. (H) Estimated numbers of granulosa cells in follicles across developmental subclasses: primordial, primary, secondary, antral, remodeling, and mitotic cycling. **Figure supplement 5**. Analysis of single-cell RNA sequencing. **Figure supplement 6**. Analysis of other cell types. **Figure supplement 7**. Hormone receptor expression and XO mouse line.

We identified up to 250 top differentially expressed genes for each group (Supplemental Dataset 5) and carried out further analysis (Metascape: Zhou et al. 2019). The development and remodeling of wave 1 follicles, and of boundary follicles giving rise to early wave 2 follicles is summarized in Figure 7.

**Figure 7.**
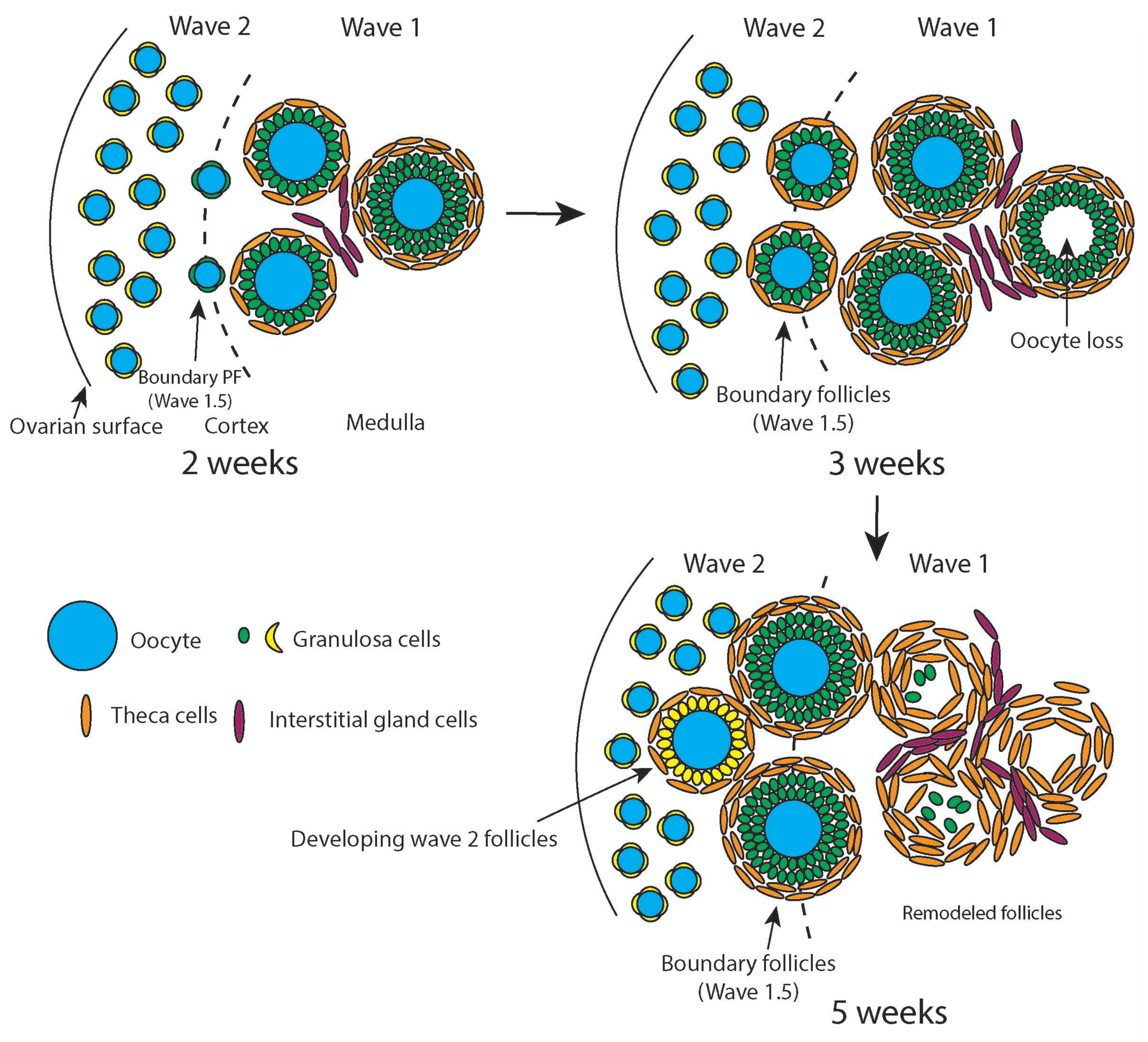
Model of wave 1 follicle development and function. Wave 1 follicles undergo remodeling during the peri-pubertal period to expand the population of androgen-producing theca and interstitial cells. Gradients between the medulla and cortex facilitate the sequential activation of primordial follicles, a process that continues into early adulthood. We refer to these sequentially activated follicles as boundary follicles (Wave 1.5). These follicles, whose granulosa cells are derived from bipotential pre-granulosa cells (BPGs), remain dormant in the medulla and cortex until recruiting signals—either positive or negative— become available. Boundary follicles are typically recruited around 2 weeks of age and constitute the majority of developing follicles by 5 weeks. In parallel, wave 1 follicles remodel to support androgen production through their remaining theca cells and associated interstitial glands.

### Comparing gene expression in wave 1 vs wave 2 follicle development

Wave 1 and wave 2 follicles begin to differ in gene expression early in fetal development, as BPGs and EPGs express different genes as early as E14.5 and diverge further by P5 (Niu and Spradling, 2020). To investigate whether gene expression differences at follicle formation persist and to look for additional wave-specific gene expression, we examined GC clusters gc2, gc7, gc10 and gc13 at 2wk, comprising all GC clusters scored as in antral follicles (Figure 5A, C). As illustrated in Figure 7, only wave 1 antral follicles are present in significant numbers at wk2 and wk3, prior to widespread wave 1 remodeling. In contrast, because of extensive wave 1 granulosa cell turnover, by wk 5 nearly all these same antral follicle clusters come from developing boundary follicles. Consequently, we compared gene expression levels during these two-time intervals for all four antral granulosa cell clusters.

The nuclear receptor *Nr5a2*, is preferentially expressed 7.8-fold higher at P5 in BPGs within growing wave 1 follicles compared to EPGs in growth-arrested wave 2 follicles (Niu and Spradling, 2020). Similarly, *Nr5a2* expression is 11-fold higher in all wk2 antral follicle cell clusters compared to primordial follicle (gc11) granulosa cells. This difference reflects the early activation of wave 1, as *Nr5a2* turns on in all wave 2 GCs as they begin to grow (Meinsohn et al. 2019). We observed that *Nr5a2* expression increases in primary and secondary follicles and reaches similar levels in wave 1 and wave 2 antral follicles (ratio = 1.02+/- 0.23). Comparable expression ratios were likewise seen between wave 1 and wave 2 antral granulosa cells for other genes that were BPG/wave 1-preferential at P5, including *Dhh* (1.04+/- 0.41), *Hsd17b1* (0.61+/- 0.14), *Ptgis* (1.08+/- 0.26), *Prlr* (1.8+/- 0.80), *Fshr* (0.89+/- 0.21) and *Igf1* (0.86+/- 0.10).

However, some genes continued to be differentially expressed between wave 1 and wave 2 follicles at the same stage of development. Wave 1 wk2 antral GCs express *Wnt4* at much higher levels (4.8 +/- 1.0) compared to Wave 2 antral CGs at either wk 5 (0.17 +/- 0.069) or wk 6 (0.14 +/- 0.070). Other genes expressed more highly in wave 1 than wave 2 antral GCs are *Igfbp5* (ratio: 27 +/- 10); *Itga6* (ratio: 6.28 +/- 0.97) and *Inha* (ratio: 1.52 +/- 0.29). Genes expressed at lower levels in wave 1 than wave 2 include: *Cald1* (ratio: 0.20 +/- 0.089), *Ghr* (ratio: 0.16 +/- 0.027), *Grb14* (ratio: 0.23 +/- 0.092), and *Mmp11* (ratio: 0.46 +/- 0.23). Thus, gene expression is not entirely equivalent in wave 1 and wave 2 follicles at corresponding stages, as has often been assumed.

Clusters gc0 and gc6 showed indications they derived from remodeling wave 1 follicle. For example, the number of gc0 and gc6 cells is low at 2 and 3 wk before substantial atresia, but jumps 9.8-fold at wk 4 (Supplemental Dataset 6). Preferentially expressed genes in gc0 and gc6 include *Cyp1b1*, *Atf3*, *Cxcl1*, *Bmp2*, *Apoe* and *Sox4*. Metascape analysis identified vasculature development (-7.7; -4.0) cell cycle (-6.7; -9.2), and positive regulation of programmed cell death (-4.1; nd), where the numbers indicate (q values log10 for gc0; gc6). These findings suggest genes and pathways that GCs on remodeling wave 1 follicles use to stimulate vasculature development, and ultimately to promote GC population turnover during wk4.

### Analyzing mesenchymal and steroidogenic gene expression

We also re-clustered the mesenchymal cells, including theca cells and stromal cells, yielding 21 new clusters to pursue a higher resolution picture of theca expansion (Figure 6E). The re-clustered mesenchymal cells fell into nine groups, late-expanding TCs (g9), early TCs (g2 and g11), *Tagln*-expressing fibroblasts (g1, g8, g14 and g17), *Col14a1*-expressing fibroblasts (g10 and g19), *Cd24a*-expressing fibroblasts (g4), *Cxcl12*-expressing fibroblasts (g3, g5 and g20), *Hoxd9*-expressing fibroblasts (g0), *Col11a1*-expressing fibroblasts (g6 and g7) and mitotic cells expressing *Mki67* (g12, g13, g15 and g18) (Figure 6F, S5D). The expression of genes in cluster16 is hard to be determined still and be named in TBD. Theca cells in g9 highly expressed steroidogenic genes such as *Star*, *Cyp11a1*, *Hsd3b1*, and *Cyp17a1* throughout wk2-wk6 and peaked in expression and cell number at w5. In contrast, TCs in cluster g11 exhibited relatively low levels of steroidogenic enzymes and stable cell numbers except at wk 5 where levels greatly increased to reach 20-50% of g9 levels. Both groups expressed LH receptor (*Lhcgr*) and prostaglandin E synthase 2 (*Ptges2*), previously implicated in TC androgen production (Erikson et al. 1985), but Lhcgr levels in g11 started lower and increased more dramatically at wk 5 than in g9.

Theca cells are not genetically capable of converting their androgen products into estrogen and usually rely on exporting them to granulosa cells for this purpose (Edstrom et al. 2007). *Star*, a crucial gene responsible for transporting cholesterol from the cytoplasm into mitochondria, is essential for steroidogenesis. The percentage of STCs expressing *Star* increased from approximately 50% to over 80% at wk 5 (Figure 6G). Our findings demonstrate that STCs increase in number and acquire a full complement of androgen biosynthetic enzymes during peri-puberty. In contrast, GCs expressing *Hsd3b1*, *Cyp19a1*, and *Hsd17a1* indicating the potential to synthesize estrogen peaked at wk 6 (Figure 6H), consistent with estrogen levels lagging androgen levels during puberty (Galas et al. 2012; Trova et al. 2021).

## DISCUSSION

### Wave 1 follicular remodeling generates a cluster of androgen-producing cells in the juvenile mouse ovary

Our work characterizes in detail the development of the first wave of mouse ovarian follicles at the cellular and gene expression levels. In agreement with earlier studies (Hirshfield and DeSanti, 1995; Gougeon, 1996; McGee and Hsueh, 2000; Eppig and Handel, 2012), 80-100% of these follicles turn over rather than generating fertilizable oocytes. We document the cellular and genetic changes within each major follicular cell type and detail the complex events that ultimately produce a large cluster of amplified steroidogenic theca cells and interstitial gland cells along with a residue of granulosa cells within the ovarian medulla by 5 wk. This process was previously termed wave 1 follicle atresia and was known to be associated with increased steroidogenesis and vascularization (Magoffin, 2005). Our analysis characterizes the cellular and molecular programs in each wave 1 follicular cell type that cause them to turn over, remodel, change in number, associate with new mesenchymal cells and to modulate their gene expression to produce steroid hormones from the ovary in a timely fashion during juvenile development. Our findings show that wave 1 follicles play a role in steroid production during juvenile development, especially at the puberty.

### Wave 1 follicles give rise to few mature oocytes that generate offspring

Boundary follicles as well as wave 1 follicles were labeled in our *Foxl2* lineage tracing experiments, and in early experiments of Zheng et al. 2014a. Boundary primordial follicles activate as early as 2 wk and produce most of the first follicles to develop fully, ovulate and generate offspring (Figure 3). It remains possible that a small number of wave 1 follicles, ∼26 or fewer, do develop to maturity. However, their contribution of wave 1 to fertility is minor at most. Thus, early activated wave 2 follicles at the medulla-cortex boundary rather than wave 1 follicles are largely responsible for early fertility. Most non-boundary wave 2 follicles only begin to activate between puberty and sexual maturity, possibly in concert with the onset of pulsatile hypothalamic-pituitary-gonadal axis activity.

### Wave 1 follicles may be a conserved feature in other animals including humans

The developing ovaries of many species produce follicles that do not complete oogenesis and ovulate (Jiménez, 2009; DeFalco and Capel, 2009). For example, an early wave of germ cells in *C. elegans* hermaphrodites develop into sperm, while during adult life only oocytes are generated (Pazdernik and Schedl, 2013). An initial wave of germ cells in the mouse testis develop during juvenile stages but break back down into individual cells, possibly to expand the stem cell pool (Lei and Spradling, 2013). Primordial germ cells in Drosophila pupal ovaries develop directly into the first ovulated oocytes, while all subsequent oocytes derive from germline stem cells that provide lifelong fertility (Zhu and Xie, 2003).

Human follicles begin developing, as in mouse, long before there is any prospect of ovulation. A fraction of primordial human follicles start to form during fetal stages, and 4-6% have progressed to the primary follicle stage prior to birth (Forabosco and Sforza, 2007). Such early developing follicles have no prospect of surviving 8 to 13 more years until the onset of puberty when they might contribute to fertility. Juvenile human ovaries 3 mo to 8 yr in age also contained 2-13% of small growing follicles that had diverged from a normal developmental pattern. 50-60% of large follicles showed signs of atresia including loss of oocytes and most granulosa cells, but retention of some theca cells (Himelstein-Braw et al., 1976). Currently, there is little understanding of the developmental mechanisms or logic underlying human fetal and juvenile follicle production. Early follicles in humans might undergo atresia and remodeling to enhance production of androgens or other steroid hormones to influence organ and brain development (Cui et al., 2013; Bell, 2018). These studies on the development of mouse wave 1 follicles and their component cell types should aid in understanding early follicular waves in many species.

### Identification of the earliest activating wave 2 follicles at the medullary-cortex boundary

A long-standing question in mammalian reproduction has been the activation order of quiescent wave 2 primordial follicles during the life cycle (McGee and Hsueh, 2000). In addition to over 250 wave 1 follicles in the medulla that begin rapid growth at birth, we identified more than 400 wave 2 primordial follicles near the medullar-cortical boundary whose GCs in 2 wk ovaries are also labeled in a mosaic fashion with *Foxl2-CreERT2* is activated at E16.5 or even earlier. Boundary follicles appear to correspond largely or entirely the primordial follicle subclass poised for early activation and located at the medulla-cortical boundary (Meinsohn et al. 2021a, b). Our data show that his substantial class of early primordial follicles do begin developing shortly after 2 wk, giving them sufficient time to become secondary and antral follicles at 5 wk. It remains possible that a small number of wave 1 follicles, ∼26 or fewer, do develop to maturity. However, it seems clear that the great majority of early progeny derive from boundary or other early developing wave 2 follicles, rather than wave 1 follicles that escape atresia.

### Mouse follicle waves express distinct genes

We found that differences in gene expression between wave 1 and wave 2 granulosa cells exist and may contribute or even control the different fates of mouse follicular waves. Previously, gene expression differences had been mapped beginning as early E14.5, and many were characterized between E16.5 to P5 (Niu and Spradling, 2020). They suggested that wave 1 follicles in the medulla differentially express several genes involved in steroid hormone production, while wave 2 granulosa cells in the cortex express genes that might predispose follicles to enter quiescence. This study extends knowledge of differences in gene expression within follicular waves during juvenile development up to 6 wk. Wave 1 follicles express much higher levels of *Wnt4* and *Igfbp5* than wave 2 follicles. Wnt4 has been genetically linked in humans to Mullerian dysgenesis and hyperandrogenism (Choussein et al. 2017) perhaps reflecting its known roles in promoting female vs male development. High *Wnt4* expression in wave 1 granulosa cells may indicate that *Wnt4*-mediated downregulation of androgen production at 2 wk becomes reduced by granulosa cell degradation, promoting the subsequent upregulation of androgen levels at puberty. *Igfbp5*, the most highly conserved of the Igfbp’s, is also a potent regulator of female reproductive tissue (Schneider et al. 2002). Thus, while changing levels of gonadotrophins in young juvenile mice likely play a role in wave 1 atresia (Edson et al., 2009), intrinsic differences in gene expression within different follicular waves at the same stage exist and have the potential to be important.

### The timing of follicular activation begins to be specified early in ovarian development

Finding that a subset of primordial follicles located near the medulla-cortex border comprise many of the earliest activated wave 2 follicles raised the question how early activation is controlled. It is well established that upregulation of insulin-mediated growth pathways accompanies follicle activation (Edson et al., 2009). Boundary follicles may become poised to active before most other wave 2 follicles because their location close to the nutrient-rich medulla enhances growth signaling, weakens quiescence and activates *Nr5a2* and *Foxl2* expression, genes which are expressed in GCs of all growing follicles. Wave 1 granulosa cells in the medulla differentially express genes promoting growth like *Igfbp5*, *Grb14* and *Ghr*, and signaling genes like *Itga6* and *Inha*, relative to wave 2 follicles growing during wks 4 and 5.

However, evidence increasingly suggests that follicular waves do not simply depend on differences in their local microenvironments. These follicle groups form at times in fetal ovary development that overlap other critical events of gonad development. The progenitors of bipotential reproductive cell types like BPGs and and mesenchymal steroidogenic precursors that arise early in both male and female gonad development remain elusive (Meinsohn, 2005; Edson et al. 2009). These cells include stem-like cells that give rise to theca cells and interstitial gland cells. We were able to preferentially mark both wave 1 follicles and boundary follicles using *Foxl2-CreERT2* as early as E14.5, suggesting that they are specified as early as bipotential and later progenitors, and involve intercellular signaling events linked to sex determination. Progress in understanding basic questions of gonad development would undoubtedly advance insight into how wave 1 and boundary follicles are specified.

### Genetic studies of wave 1 follicles

We explored genotypes that reduce the number of wave 1 follicles in order to look for the genetic consequences of decreased wave 1 activity. In Turner syndrome, wave 1 follicle development is reported to be reduced significantly (Miura et al. 2017). This imbalance might decrease androgen levels, potentially affecting juvenile development. Consistent with hormonal disruption, hormone replacement therapy significantly improves the development of individuals with Turner syndrome (Klein et al., 2018). Moreover, androgens are known to play an important role in this effect (Zuckerman-Levin et al., 2009; Viuff et al., 2022).

We examined mouse XO mice and looked for changes in wave 1 follicle numbers (Figure S7B-C). In our experiments, wave 1 follicle numbers were reduced relative to levels in XX control females at wk1 of development. However, the reduction was transient and by 2 wk, before remodeling and theca cell expansion, XO and XX females had regained a similar number of wave 1 follicles. Consequently, we could not draw any conclusions regarding the functional role of wave 1 follicles and their remodeling from these studies.

### Steroid hormones mediate sexual development in juvenile mice and during peri-puberty

Steroid hormones influence the development of sexually dimorphic organs including the brain (review Simerly, 2002; Arnold, 2009). They have multiple sources, including steroids in the maternal circulation during pregnancy, as well as subsequent systemic and local steroid production during juvenile stages. Wave 1-derived theca cell numbers are high during wk 4-5 of juvenile development. a period known as peri-puberty that has been associated with sexual behavior development and with high levels of androgens, as well as relatively low estrogen production (François et al. 2017; Trova et al. 2021; Devillers et al., 2023). While the sensitivity of the brain to steroid influences is probably declining during this period, sexually dimorphic circuits are still being shaped in a manner that is important for sexually dimorphic behaviors in female and male rodents (Sisk and Zehr, 2005; Templin et al., 2019; Trova et al., 2021; Yoest et al., 2023). Potential targets of this influence include the anteroventral periventricular nucleus (AVPV) which is larger in female than male rodents and helps regulate female reproductive functions such as ovulation and gonadotrophin production. Sex hormones produced in females during puberty also enhance cell death in the primary visual cortex, reducing its volume in females compared to males. Thus, wave 1 generated steroid production may influence the sexual development.

### Wave 1 follicle remodeling, polycystic ovarian syndrome and human follicular waves

The work reported here may be relevant to the origin and high frequency of polycystic ovary syndrome (PCOS) (McCartney and Marshall, 2016). A possible connection between follicular atresia, androgen excess and PCOS has been discussed previously (reviewed in Erikson et al., 1985; Magoffin, 2005). Interestingly, some histological characteristics of remodeling mouse wave 1 follicles resemble follicles within the ovaries of individuals with PCOS (Chang, 2007; Franks and Hardy, 2010; Walters et al., 2019). A good place to start toward further progress on this connection would be to characterize the development and gene expression of early human follicles and compare them to the findings reported in other systems, including mice.

## MATERIALS AND METHODS

### Antibodies

| Name | Source | Identifier | Dilution |
| --- | --- | --- | --- |
| Anti-EGFP (Chicken) | Aves labs | GFP-1020 | 1:300 |
| Anti-DDX4 (Rabbit) | Abcam | ab13840 | 1:300 |
| Anti-MSY2 (Mouse) | Abcam | sc-393840 | 1:100 |
| Anti-tdTomato (Goat) | Origene Tech | AB8181-200 | 1:300 |
| Anti-HSD3B1 (Mouse) | Santa Cruz | sc-515120 | 1:100 |
| Anti-CYP17A1 (Rabbit) | Cell Signaling Technology | 94004T | 1:100 |
| Anti-PLIN1 (Rabbit) | abcam | ab3526 | 1:100 |
| DAPI | Sigma | D9542-10mg | 0.5ug/ml |
| Alexa Fluor® 488 AffiniPure™ Donkey Anti-Chicken IgY (IgG) (H+L) | Jacksonimmuno | 703-545-155 | 1:300 |
| Alexa Fluor® 594 AffiniPure™ Donkey Anti-Goat IgG (H+L) | Jacksonimmuno | 705-585-147 | 1:300 |
| Alexa Fluor 488 donkey anti-mouse IgG | invitrogen | A21202 | 1:300 |
| Alexa Fluor 568 donkey anti-mouse IgG | invitrogen | A10037 | 1:300 |
| Alexa Fluor 488 donkey anti-rabbit IgG | invitrogen | A21206 | 1:300 |
| Alexa Fluor 568 donkey anti-rabbit IgG | invitrogen | A10042 | 1:300 |
| Alexa Fluor 647 donkey anti-rabbit IgG | invitrogen | A31573 | 1:300 |

### Primers

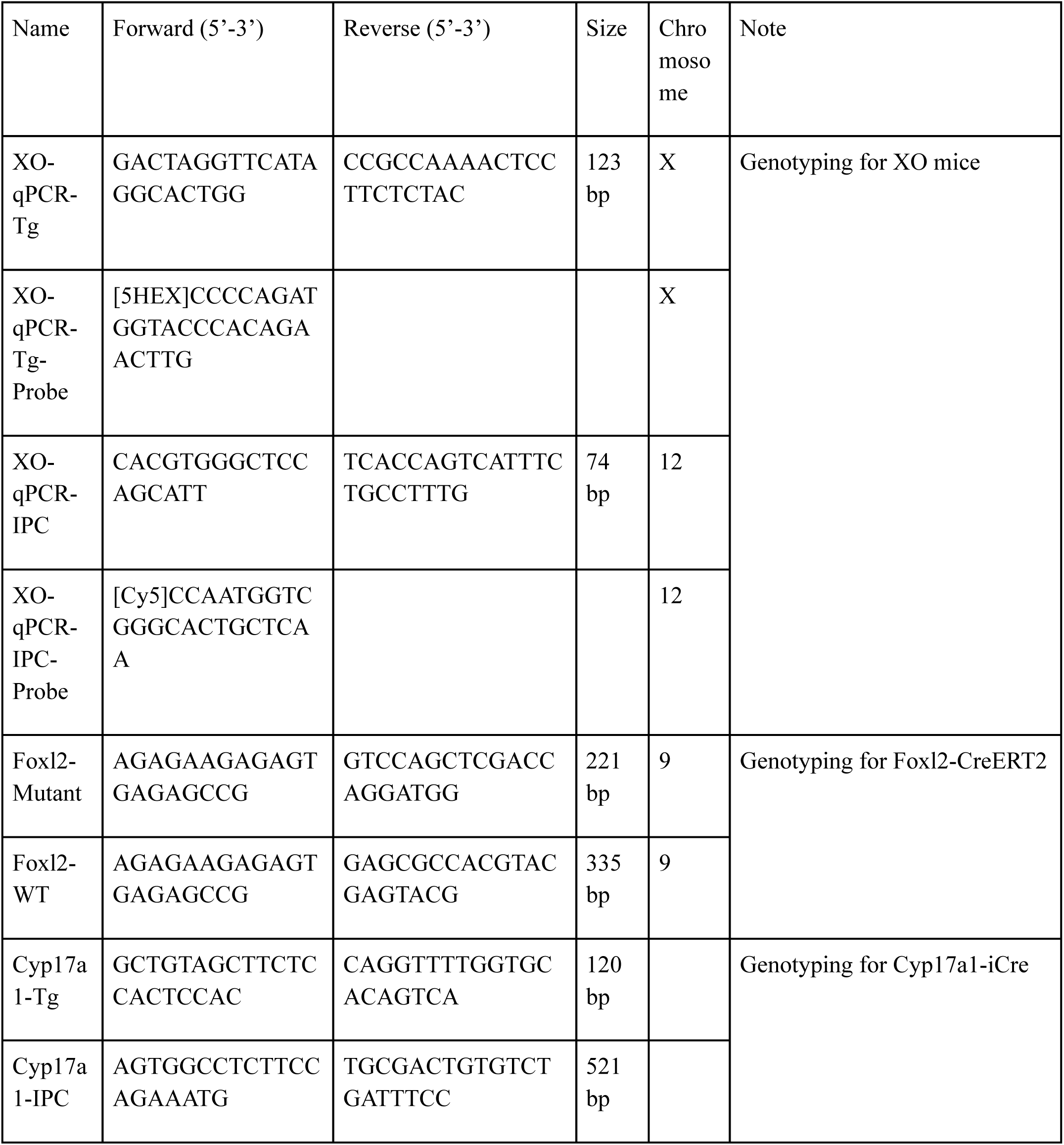

### Experimental mice

All animal experiments were conducted in compliance with the Institutional Animal Care and Use Committee guidelines of the Carnegie Institution of Washington (CIW). The mice were housed in specific pathogen-free facilities at CIW. The strains of mice used in the study were acquired from Jackson Laboratory, including C57BL/6J (B6, strain #: 000664), DBA/2J (DBA2, strain #: 000671), B6;129P2-*Foxl2^tm1(GFP/cre/ERT2)Pzg^*/J (*Foxl2-CreERT2*, strain #: 015854), STOCK XO/J (XO, strain #: 036414), B6CBACaF1/J-*A^w-J^/A* (Strain #:001201) and B6;SJL-Tg(Cyp17a1-icre)AJako/J (*Cyp17a1-icre*, strain #: 028547). The *R26R-LSL-EYFP* reporter mice B6.129X1-*Gt(ROSA)26Sor^tm1(EYFP)Cos^*/J (*R26R-LSL-EYFP*, strain #: 006148) were acquired from Jackson Laboratory and backcrossed onto 129S1/SvImJ (129S1, strain #: 002448) background in our lab. The R26R-LSL-tdTomato reporter mice were generously provided by Chen-Ming Fan at the CIW. CD-1 IGS Mouse (strain #: 022) was obtained from Charles River Laboratories.

### Lineage tracing experiments

To visualize the expression patterns of *Foxl2* and *Cyp17a1*, we generated reporter mouse lines by crossing *R26R-EYFP* or *R26R-tdTomato* mice with *Foxl2-CreERT2* or *Cyp17a1-iCre* (Shimshek et al., 2002; Zhou et al., 2022) mice. The presence of a vaginal plug was designated as embryonic day 0.5 (E0.5). Pregnant females were injected intraperitoneally (IP) with 1 mg tamoxifen (TAM) (MilliporeSigma, #T5648, 10 mg/ml dissolved in corn oil) per 35 g body weight at E16.5 or E14.5 (*Foxl2-CreERT2*). Pups were either delivered naturally or, if not born on the expected delivery day, delivered by C-section and fostered by a CD1 female mouse. The day of birth was considered postnatal day 0 (P0). We collected gonads from fetuses or ovaries from pups at the indicated stages for analysis. All experiments were conducted with at least three biological replicates from two litters and were repeated twice. The loss of one *Foxl2* allele in *Foxl2-CreERT2* mice, along with tamoxifen administration, does not significantly impact ovarian development or overall mouse health.

To investigate the expression of *Foxl2* in remodeled follicles, tamoxifen was injected IP at a concentration of 1 mg per 35 g weight into female mice at 5 weeks of age. The ovaries were dissected one week later and subjected to whole mount staining for analysis.

### Immunostaining

Whole-mount staining was performed as previously described (Li et al., 2017). Briefly, gonads or ovaries were fixed in 4% paraformaldehyde (PFA) overnight at 4°C. The tissues were then washed three times in a washing buffer (0.3% Triton X-100 and 0.5% thioglycerol in PBS) and incubated in blocking buffer (1% normal donkey serum, 1% BSA, and 0.3% Triton X-100 in PBS) overnight at 37°C. The samples were incubated with primary antibody in dilution buffer (1% BSA and 0.3% Triton X-100 in PBS) for two days at 37°C. After washing in the washing buffer overnight at 37°C, the samples were incubated with secondary antibodies in dilution buffer for another two days at 37°C, followed by washing in the washing buffer with DAPI overnight at 37°C. To facilitate whole-tissue imaging, the clearing buffer Ce3D (prepared by dissolving 4 g Histodenz in 2.75 ml of 40% [v/v] N-methylacetamide) was applied for an additional day. The samples were imaged using a Leica TCS SP8 confocal microscope or a Leica Stellaris 8 DIVE confocal microscope. Images were analyzed using Fiji or Imaris.

For fluorescence cryosection staining, the dissected tissues were fixed in 4% PFA overnight at 4°C. After washing three times in PBS, the tissues were dehydrated sequentially in 10% and 30% sucrose solutions and embedded in O.C.T. for cryosectioning. The slides were sectioned at a thickness of 5 μm and stored at -20°C until staining. After antigen retrieval in 0.01% sodium citrate buffer at boiling temperature for 20 minutes, the slides were sequentially blocked for 1 hour at room temperature, incubated with primary antibodies overnight at 4°C, and then with secondary antibodies for 1 hour at room temperature. This was followed by counterstaining with DAPI for 10 minutes before fixing in mounting medium. The samples were then imaged with an upright SP5 microscope, and the images were analyzed using Fiji.

### Oil Red O Staining

Oil Red O Staining was performed using Oil Red O stain kit (Abcam, ab150678) following the manufacturer’s protocol. Briefly, the slides were placed in propylene glycol for 5 minutes at room temperature, followed by incubation in Oil Red O solution at 60°C for 10 minutes. The slides were then differentiated in 85% propylene glycol for 1 minute, followed by rinsing twice in distilled water. The slides were then incubated in hematoxylin for 1 minute, rinsed thoroughly in tap water and twice in distilled water. The slides were ready for imaging after applying a coverslip using an aqueous mounting medium. Imaging was done using a Nikon Eclipse E800 microscope.

### Genotyping

The tissue was dissected and incubated in a lysis buffer (30 mM Tris-HCl, 0.5% Triton X-100, 200 μg/ml Proteinase K) at 55°C overnight in a water bath. The lysis buffer was then inactivated by heating at 95°C for 5 minutes.

For genotyping using conventional PCR, 1 μl of the sample was added to the PCR mixture (primer sequences are provided in the supplemental data). For quantitative PCR (qPCR) to genotype XO mice, primers and probes (sequences included in the supplemental data) were designed to amplify and detect target DNA sequences from both the X chromosome and an autosome. qPCR was performed using a CFX 96 Real-Time PCR machine. The amplification cycle threshold for the X chromosome was compared to that of the autosome to determine the genotype, distinguishing between XO (one X chromosome) and XX (two X chromosomes) females.

### Single cell RNA sequencing

Tissue samples were first dissected in PBS and dissociated using 0.25% Trypsin/EDTA (Thermo Fisher, #25200056) at 37°C for approximately 20 minutes. Dissociation was halted by adding FBS (Thermo Fisher, #10439016) to a final concentration of 10%. The resulting cell suspension was filtered through a 100 μm strainer (Fisher, #22-363-549) to remove any remaining undissociated structures. Cells were pelleted by centrifugation using an Allegra X-14R Centrifuge (Beckman Coulter) at 300 × g for 3 minutes and suspended in PBS containing 0.04% BSA (PBSB). To further eliminate cell clumps, the samples were filtered through a 40 μm strainer (Fisher, #22-363-547), followed by centrifugation at 300 × g for 5 minutes. The cells were then resuspended in PBSB and subjected to FACS sorting (BD FACSAria™ III Cell Sorter) to collect live cells, with SYTOX™ Orange Dead Cell Stain (Thermo Fisher, S34861) used for viability selection. After a final centrifugation at 300 × g for 5 minutes, the cells were resuspended in appropriate volume of PBSB to achieve a concentration of 1000 cells/μl before proceeding with single-cell sequencing.

Libraries of the single cells were prepared using the Chromium Next GEM Single Cell 3’ Reagent Kits v3.1(PN-1000128) from 10X Genomics, with the goal of capturing 10,000 cells per sample. Sequencing was performed using the NextSeq 500 platform from Illumina according to 10X Genomics recommendations: paired-end, single index reads with Read 1 being 28 cycles, Index 1 being 8 cycles, and Read 2 being 91 cycles. Sequencing was performed to a depth of 50,000 reads per cell. The single cell sequencing datasets were initially processed using the Cell Ranger pipeline (v3.1.0 and v6.0.1) with default settings.

Further analysis of the scRNA-seq data was performed using the Seurat package (v4.1.0). The count data were read and transformed into a Seurat object using the ‘Read10X’ and ‘CreateSeuratObject’ functions. The libraries (2 weeks, 3 weeks, 4 weeks, 5 weeks, and 6 weeks) were preprocessed with ‘NormalizeData()’, followed by ‘FindVariableFeatures()’, ‘FindIntegrationAnchors()’, and ‘IntegrateData()’ using default settings. The integrated object was further processed using ‘ScaleData()’, ‘RunPCA(npcs = 50)’, ‘RunUMAP(dims = 1:30)’, ‘FindNeighbors(dims = 1:30)’, ‘FindClusters(resolution = 1.0)’, and ‘RunTSNE(dims = 1:30)’ with other default settings to determine cell clusters. Further clustering of granulosa cells and mesenchymal cells was performed using ‘subset ()’, followed by the cell clustering methods mentioned above.

### The electron microscope

Mice ovaries were excised and fixed in a solution of 2% PFA and 2.5% glutaraldehyde in 0.1 M PIPES buffer (MilliporeSigma, #P6757, pH 7.4) at 4°C overnight. The ovaries were then washed and treated with 50 mM glycine in 0.1 M PIPES buffer for 15 minutes, followed by washing with 0.1 M PIPES buffer. Tissue pieces were then postfixed with 1% osmium tetroxide and 1.5% potassium ferrocyanide in 0.1 M PIPES buffer for 60 minutes, washed with water, and stained en bloc with 1% (w/v) uranyl acetate for 60 minutes. The ovaries were then serially dehydrated using graded ethanol solutions: 30%, 50%, 75%, 85%, 95%, and 100% ethanol, followed by two exchanges of 100% acetone. After dehydration, the ovaries were infiltrated and embedded in EMbed 812 Epoxy resin (Electron Microscopy Sciences) according to the manufacturer’s recommendations. Ultrathin sections at about 70 nm thick were cut on a Leica EM UC7 ultramicrotome and examined using a Hitachi HT7800 transmission electron microscope operated at 80 kV. Images were acquired with an AMT NanoSprint 12 camera.

### Statistical analysis

The data are reported as means ± SD of biological replicates, and no statistical methods were used to determine the sample size. Differences between two groups were compared using a two-tailed Student’s *t*-test. A *p*-value of <0.05 was considered statistically significant and marked with one star (*), *p* < 0.01 with two stars (**), *p* < 0.001 with three stars (***), and *p* < 0.0001 with four stars (****).

**Table 1.** Summary of whole mount C57BL/6J ovary analyses.

| Postnatal age | Total follicles/<br>ovary | Antral | Wave 1<br>follicles/ovary | Reference files | Ovaries<br>analyzed |
| --- | --- | --- | --- | --- | --- |
| P5-7 | 3822 +/- 838 | 0 | 278 +/- 17 | 1wk_1.mp4;<br>1wk_2.mp4 | 4 |
| P10-P14 | 3304 +/- 926 | 0 | 293 +/- 36 | 2wk_1.mp4;<br>2wk_2.mp4 | 3 |
| 3 wk | 3089 +/- 669 | 23 +/- 2 | ND | 3wk_1.mp4<br>3wk_2.mp4 | 3 |
| 4 wk | ND | 35 +/- 2 | ND | 4wk_1.mp4<br>4wk_2.mp4 | 4 |
| 5 wk | ND | 24 +/- 4 | ND | 5wk_1.mp4<br>5wk_2.mp4 | 5 |

Numbers are mean and standard deviation.

## Data availability

The scRNAseq data have been deposited at the NIH GEO database under accession number GSE268466.

## Funding

Howard Hughes Medical Institute

## Supplementary Materials

Figs. S1 to S7

Movies S1 to S10

Datasets S1-S6

## ACKNOWLEDGEMENTS

We are grateful to Allison Pinder and Dr. Fredrick Tan for assisting with the genomic experiments and analyses. We thank Dr. Mamhud Siddiqi for his help with optical microscopy and image analyses. We thank Dr. Ru-Ching Hsia for expert electron microscopy. We thank the Carnegie Embryology support staff and Spradling lab members for comments on the study, especially Haolong Zhu for assistance with scRNA-seq data analysis.

## Author contributions

Investigation: QY, Supervision and Funding: ACS, all other: QY and ACS.

## Competing interests

None.

## Supplemental Figure Legends

**Figure supplement 1.**
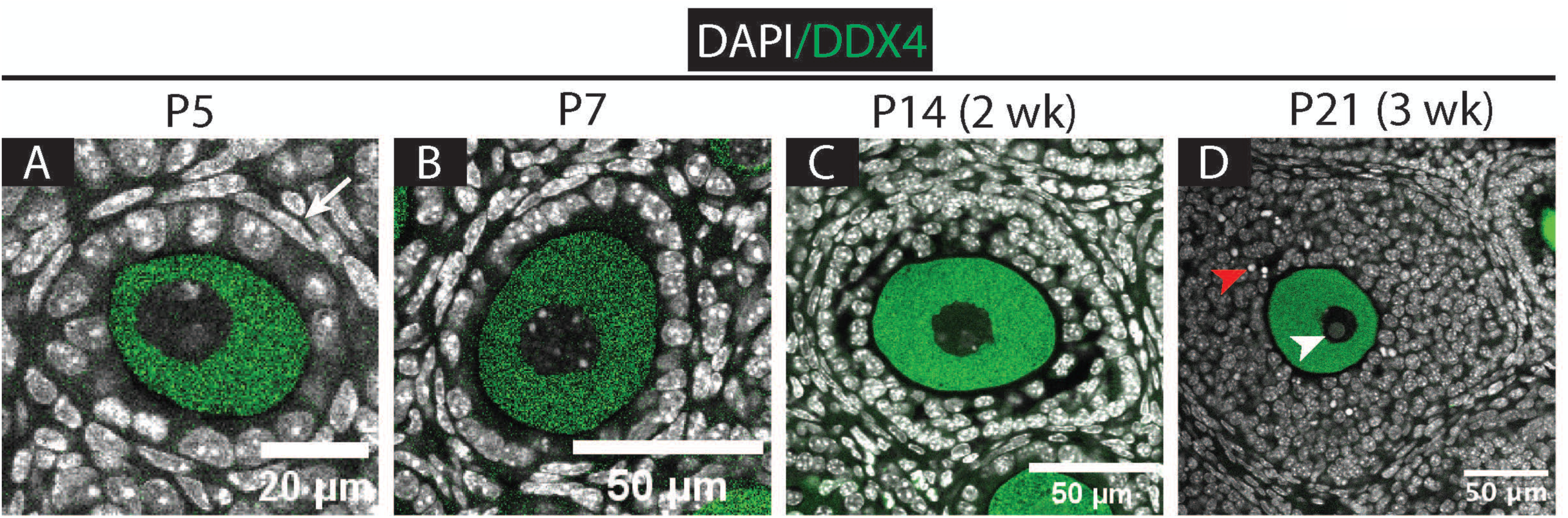
Wave 1 follicle development in the early juvenile ovary. (**A-D**) Representative images of wave 1 follicles stained for DDX4 (green) at P5, P7, P14 (2 wk), and P21 (3 wk). A: Early primary follicle with theca cells (arrow); B: Late primary follicle; C: Secondary follicle; D: Antral follicle. White arrowhead: non-surrounded nucleus; Red arrowhead: Pyknotic nuclei of granulosa cells. Scale bars in, A: 20 µm; others: 50 µm.

**Figure supplement 2.**
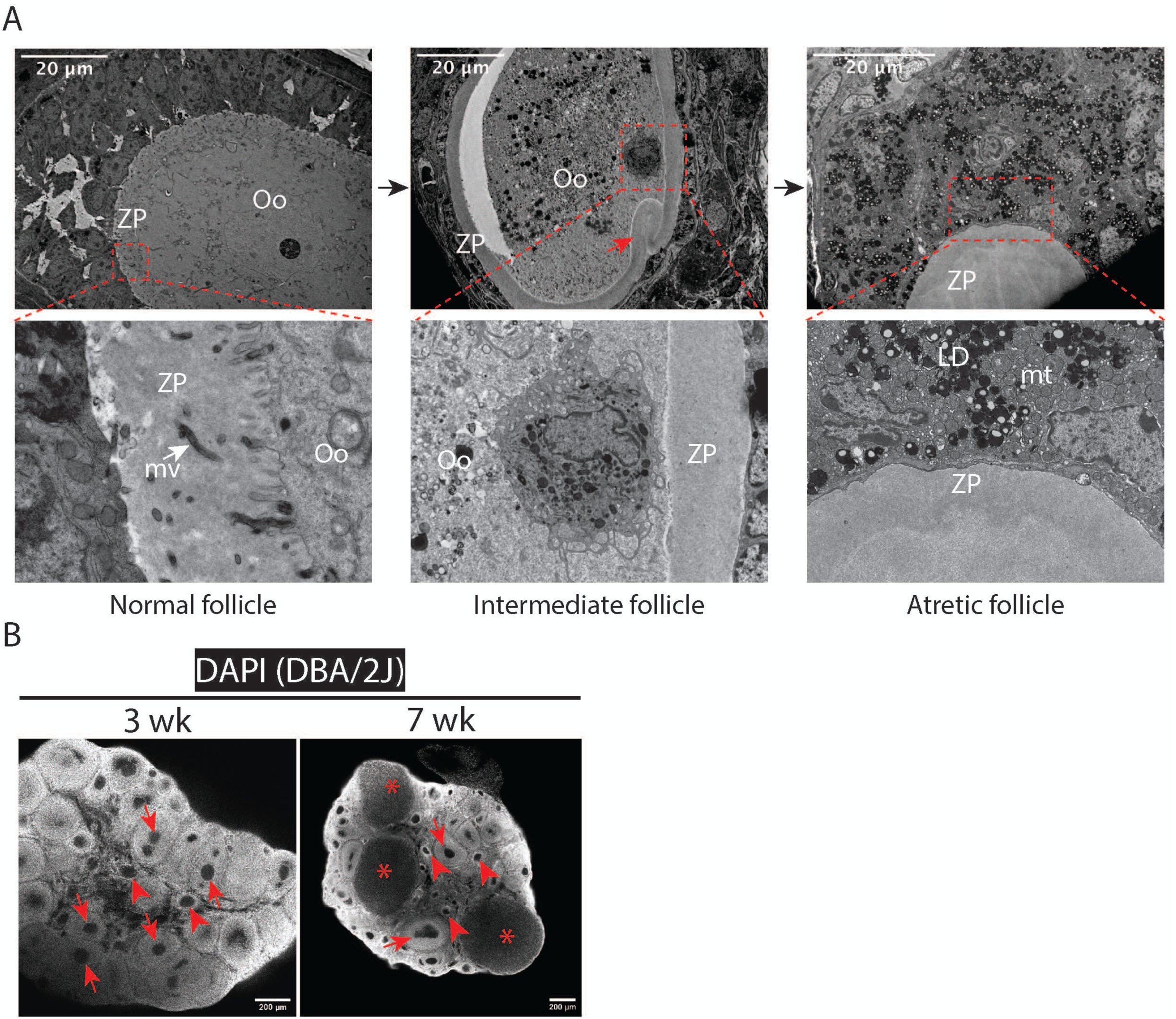
Wave 1 follicles begin to remodel by 3 weeks. (**A**) Electron microscopy of follicles at 5 wk ovary. Left: Normal follicle. Middle: intermediate follicle, in which the oocyte cytoplasm shrinks and begins degrade, indicated by the space between the oocyte and ZP, the absence of microvilli, and folding of the ZP (red arrow). Right: Remodeled follicle, where the oocyte has disappeared but the ZP remains. Surrounding theca and interstitial gland cells further accumulate lipid droplets. The boxed region is magnified in the lower panel. Abbreviations: Oo = oocyte; ZP = zona pellucida; LD = lipid droplet; mt = mitochondria; mv = microvilli. Scale bars = 20 µm. (**B**) DAPI-stained DBA/2J ovaries showing remodeling wave 1 follicles (arrowheads) and normally developing follicles (arrows). Left (3 wk): Early remodeling follicles are observed in the central medulla (arrowheads). Right (7 wk): Wave 1 follicle derivatives retain an empty shape (arrowheads), remaining visible alongside normal follicles (arrows) and corpora lutea (asterisks). Ovaries were stained and imaged as whole mounts; only a single section is shown here. Scale bars = 200 µm.

**Figure supplement 3.**
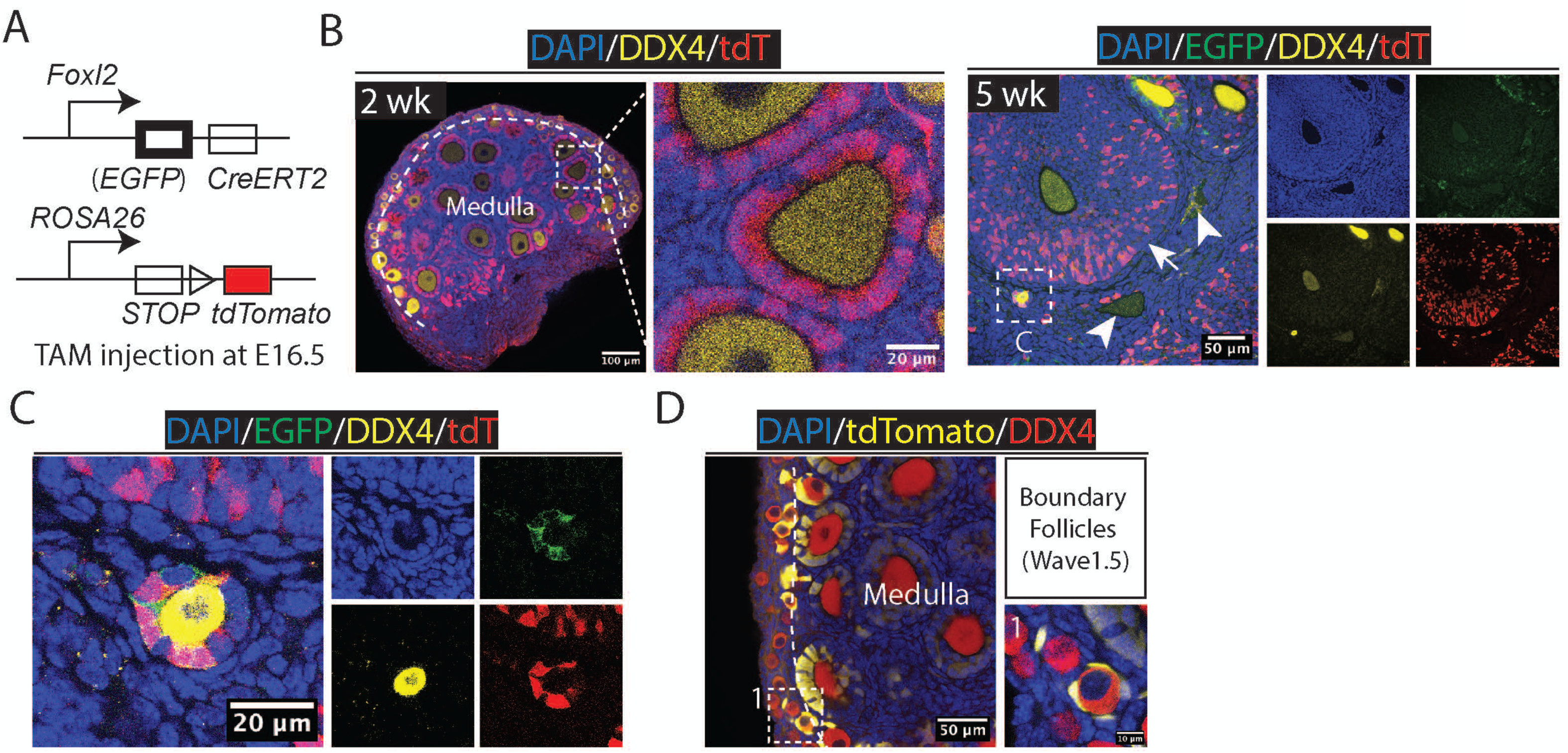
Lineage labeling using *Foxl2-CreERT2* and *R26R-LSL-tdTomato*. (**A**) Schematic diagram of the experiment. The *Foxl2-CreERT2* mouse line was generated with EGFP inserted into the *Foxl2* locus, resulting in weak EGFP signal detectable in the cytoplasm using an EGFP antibody. To rule out the possibility that the observed signal originated from the *Foxl2* locus rather than *ROSA26-EYFP*, we repeated the experiment using the *Foxl2-CreERT2; R26R-LSL-tdTomato* mouse line. (**B**) At 2 wk, wave 1 follicle are labeled. By 5 wk, ome have turned over, as indicated by cavities left by disappeared oocytes and remaining tdTomato signals (arrowheads), while others have developed into antral follicles (arrow). One developing early primary follicle in the boxed region is shown in panel C. Scale bars are included in the images. (**C**) The EGFP channel shows a weak cytoplasmic signal from the Foxl2 locus, distinct from the strong, evenly distributed expression of EYFP from the *ROSA-EYFP* reporter shown in Figure 3/4. Scale bar, 20 µm. (**D**) Boundary follicles (enlarged on the right) were observed in the 2 wk ovary of *Foxl2-CreERT2; R26R-LSL-tdTomato* mouse line injected with TAM at E16.5. Scale bars: left, 50 µm; right: 10 µm.

**Figure supplement 4.**
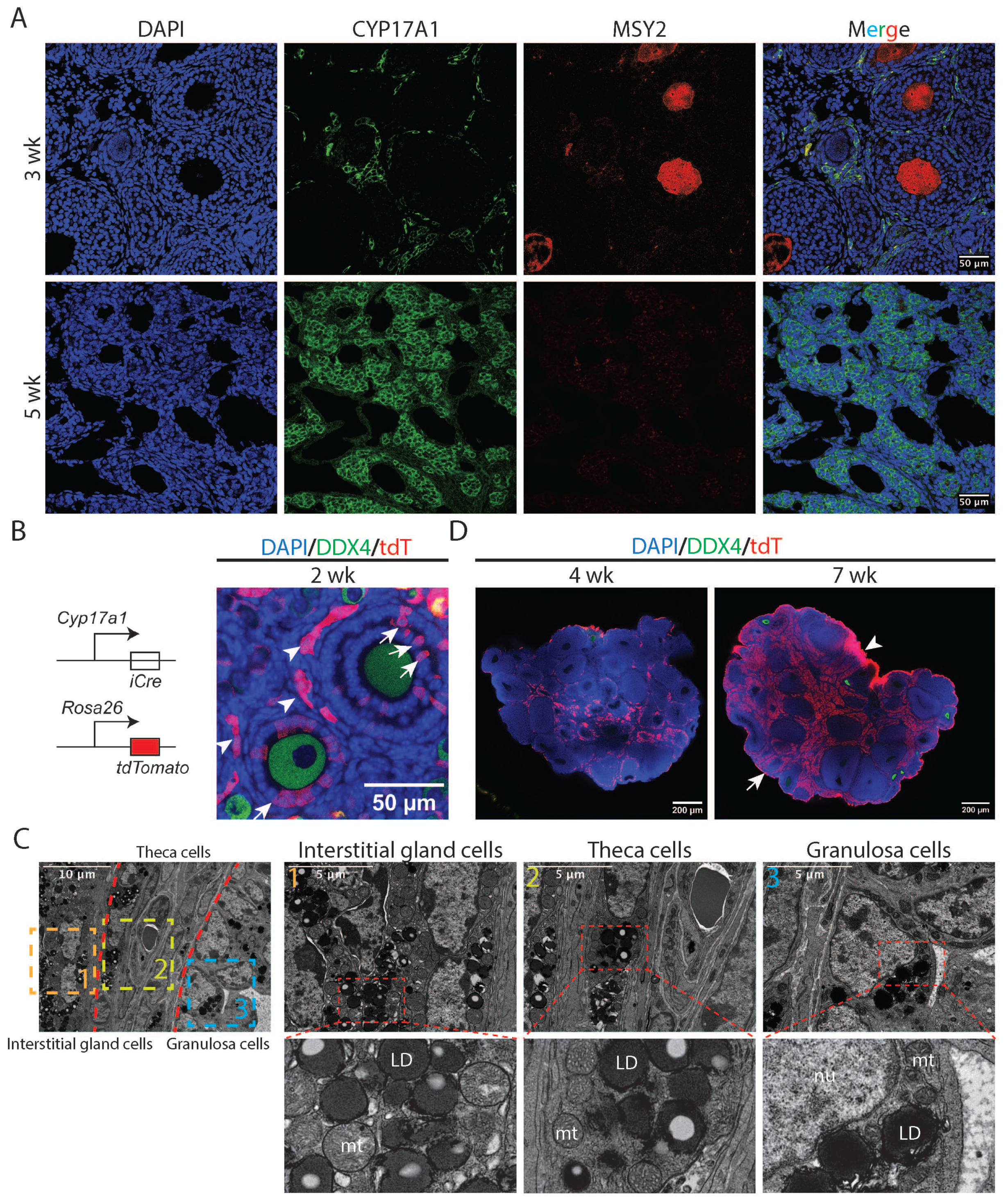
Expansion of theca cells and its related cells - interstitial gland population. (**A**) Immunofluorescence staining of CYP17A1 protein in the ovary shows a significant increase of theca cells and interstitial gland within the stroma from 3 wk to 5 wwk. Scale bars = 50 µm. (**B**) Immunofluorescence staining of *Cyp17a1* lineage-traced ovary at 2 wk shows initial expression in theca cells and associated cells resembling interstitial gland cells (arrowheads). Additionally, a few granulosa cells were labeled (arrows), as previously reported (Bridges et al., 2008). Scale bar = 50 µm. (**C**) Structurally, steroidogenic theca cells closely resemble interstitial gland cells. In the electron microscopy images, red lines delineate distinct zones enriched in interstitial gland cells (1), theca cells (2), and granulosa cells (3), with each region magnified below. Zone 1 (rectangle labeled “1”) represents interstitial gland cells. Zone 2 (rectangle labeled “2”) contains steroidogenic theca cells. Zone 3 (rectangle labeled “3”) corresponds to granulosa cells. Abbreviations: nu – nucleus; LD – lipid droplet; mt – mitochondria. Scale bars are included in the images. (**D**) *Cyp17a1* lineage-traced ovary at 4 wk and 7 wk shows a marked increase in the number of labeled theca cells and interstitial gland cells by 7 weeks. Extensive signal is also observed in the rete ovarii (arrowhead), indicating Cyp17a1 expression, as well as in the ovarian epithelium (arrow). Scale bars = 200 µm.

**Figure supplement 5.**
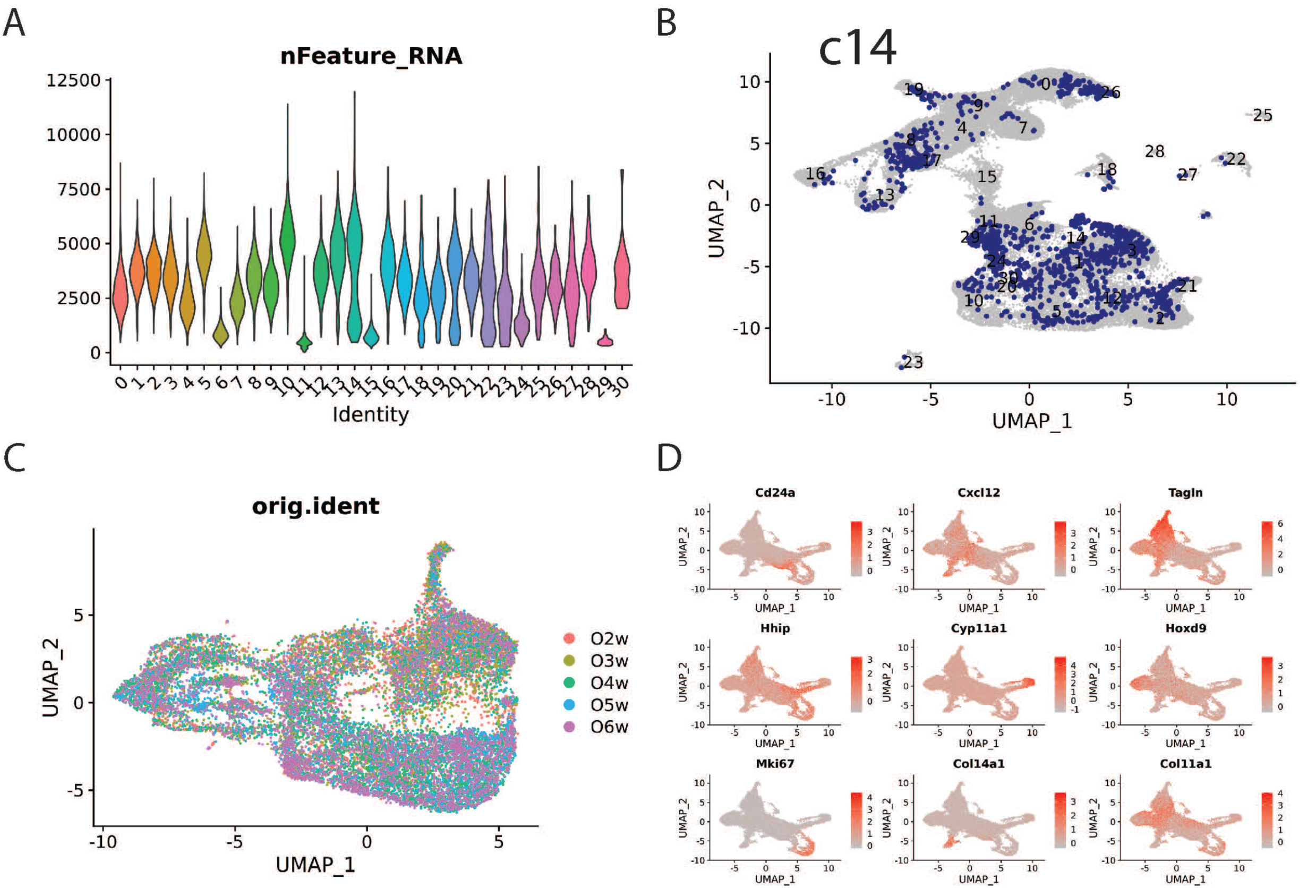
Analysis of single-cell RNA sequencing. (**A**) nFeature_RNA of the scRNA-seq data shows that clusers 6, 11, 15, 29 have low UMI. (**B**) Highlighting cluster 14 reveals its dispersed distribution. (**C**) The distribution of samples across clusters shows that cells from each sample contribute to all clusters (orig.ident). (**D**) Feature plot of marker gene expression from the reanalysis of mesenchymal cells, including late theca cells (*Cyp11a1*), early theca cells (*Hhip*), Tagln+ fibroblasts (*Tagln*), Hox9+ fibroblasts (*Hox9*), Col11a1+ fibroblasts (*Col11a1*), Cxcl12+ fibroblast (*Cxcl12*), Col14a1+ fibroblasts (*Col14a1*), Cd24a+ fibroblasts (*Cd24a*) and mitotic cells (*Mki67*).

**Figure supplement 6.**
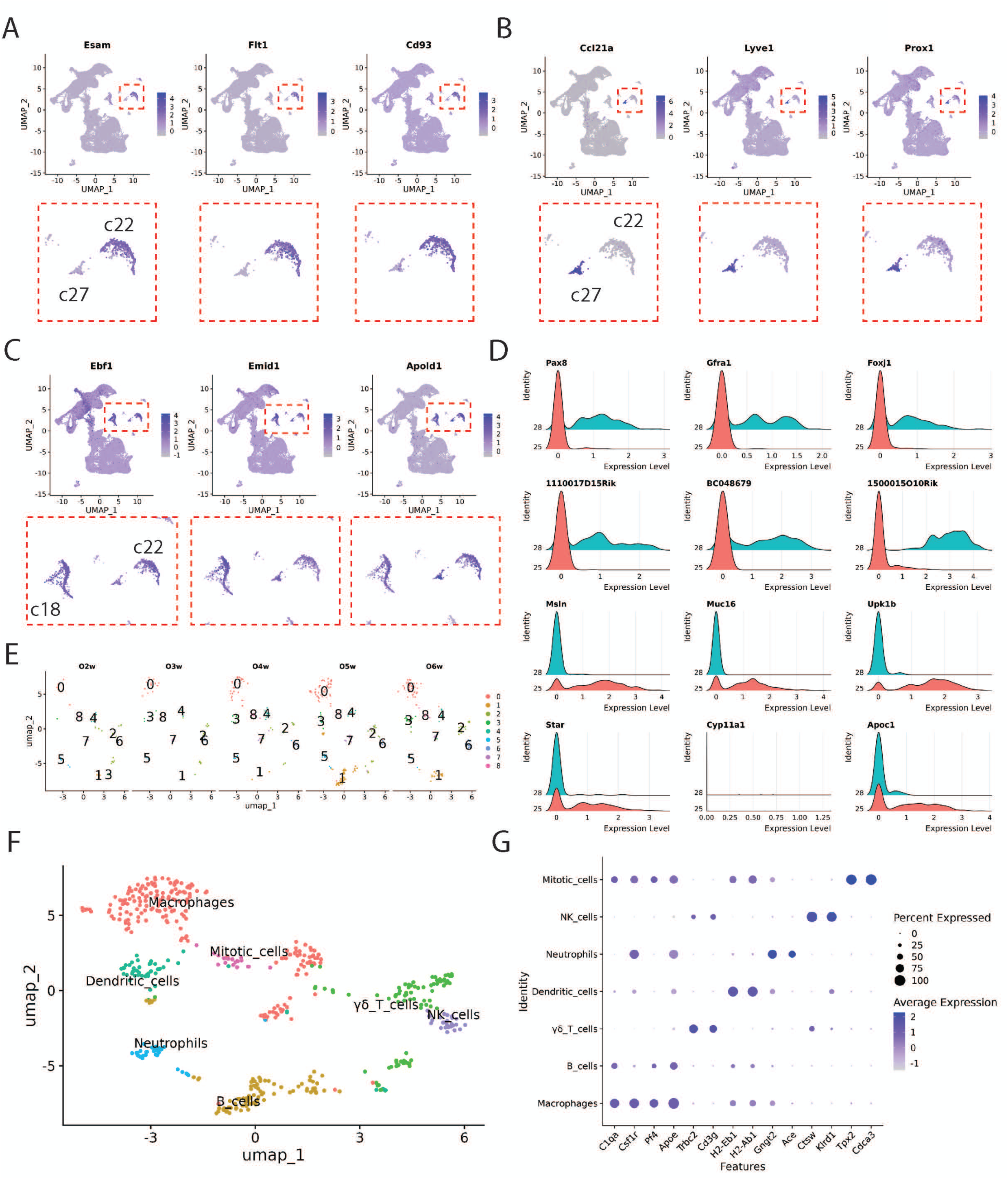
Analysis of other cell types. (**A**) Expression plot of representative genes highly expressed in cluster 22, identified as endothelial cells in blood vessels, including *Esam*, *Flt1*, and *Cd93*. Red boxes in the top panel are enlarged in the bottom panel. cluster 27: Endothelial cells from lymphatic vessels. (**B**) Expression plot of representative genes highly expressed in cluster 27, identified as endothelial cells in lymphatic vessels, including *Ccl21a*, *Lyve1*, and *Prox1*. Red boxes in the top panel are enlarged in the bottom panel. (**C**) Expression plot of representative genes shared between pericytes and endothelial cells, including *Ebf1*, *Emid1*, and *Apold1*. Red boxes in the top panel are enlarged in the bottom panel. (**D**) Ridge plot of representative gene expression in epithelial cells from the ovarian surface epithelium (cluster 25) and the rete ovarii (cluster 28). (**E**) Further analysis of hematopoietic cells using re-clustering in Seurat identified 9 clusters across five samples. (**F**) The deduced hematopoietic cell subgroups are labeled and indicated by colored regions, including macrophages (clusters 0, 4, and 7), dendritic cells (cluster 3), neutrophils (cluster 5), B cells (cluster 1), T cells (cluster 2), NK cells (cluster 6), and mitotic cells (cluster 8). (**G**) Dot plot showing marker gene expression across different hematopoietic cell types.

**Figure supplement 7.**
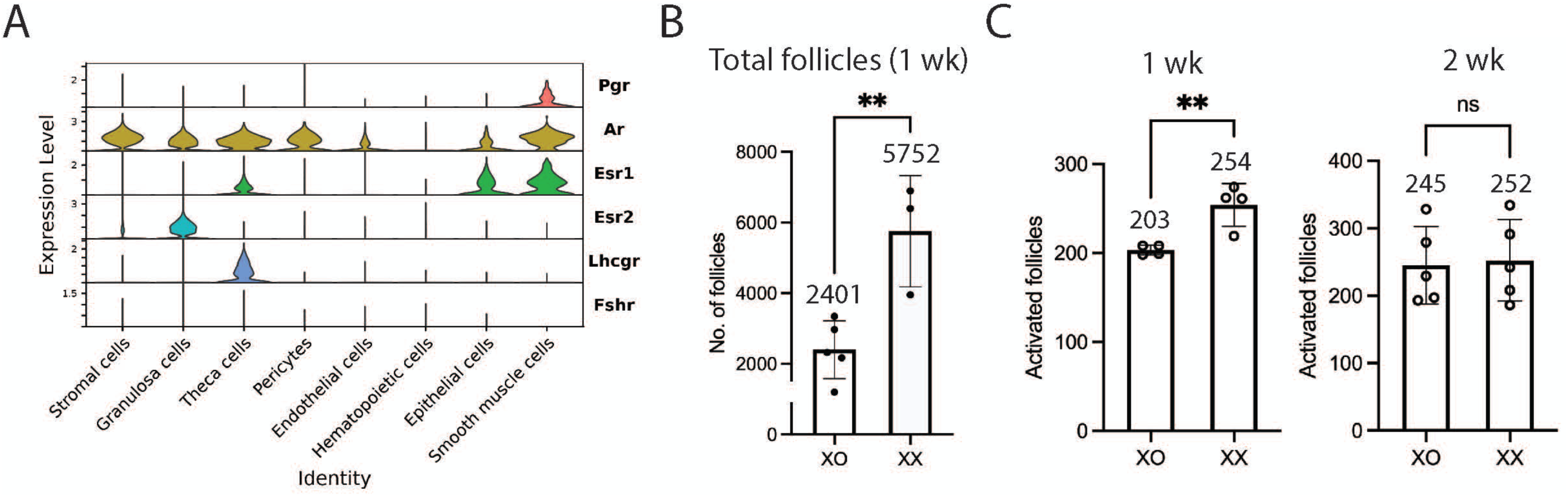
Hormone receptor expression and XO mouse line. (**A**) Expression levels of hormone receptors (Y-axis) across different cell clusters (X-axis). Androgen receptor (*Ar*) is expressed in nearly all cell types except hematopoietic cells. (*Esr1*) is primarily expressed in theca cells, epithelial cells, and smooth muscle cells, whereas estrogen receptor 2 (*Esr2*) is expressed in granulosa cells as expected. No expression of follicle-stimulating hormone receptor (*Fshr*) is detected in any cell type, while luteinizing hormone/choriogonadotropin receptor (*Lhcgr*) is specifically expressed in theca cells. Progesterone receptor (*Pgr*) is present in smooth muscle cells but absent in granulosa cells. (**B**) The total follicle count in XO mice at 1 week is significantly lower compared to control XX mice. Follicle counts were performed using Imaris software on whole-mount stained ovaries. N = 3–5. ** *p* < 0.01. (**C**) By 2 weeks, statistical difference between XO and XX mice were observed. Follicle counts were performed using Imaris software on whole-mount stained ovaries. N = 4–5. ** *p* < 0.01; ns, not significant.

## Notes

### Competing Interest Statement

The authors have declared no competing interest.

### Summary of Updates

Improvements were made to the figures, especially by swapping some panels previously in the Supplemental figures to Figures 1 and 3. An higher quality file of the same material was swapped for Figure 2C and 2D and the relationship of data figures and their summary graphs was improved. Small changes to the labeling were made to improve clarity in Figures 4-7. The entire manuscript was edited to improve clarity. The tradition term "atresia" to used describe the changes in wave 1 follicles initially, before explaining that these changes constituted a form of remodeling.

